# Stromal innervation is prognostic in non-muscle-invasive bladder cancer and predicted by a distinct microenvironmental transcriptomic signature

**DOI:** 10.64898/2026.09.21.751588

**Authors:** Cailin S Deiter, Florus C de Jong, Mitchell Olislagers, Kimberly R Jordan, Tahlita C M Zuiverloon, James C Costello

**Author notes:** Corresponding Author James C Costello, University of Colorado Anschutz Department of Pharmacology Mail Stop 8303, RC1-South, 12801 East 17th Ave Aurora CO 80045.

## Abstract

Peripheral nerves are increasingly recognized as active components of the tumor microenvironment (TME). Yet, their role in non-muscle-invasive bladder cancer (NMIBC) and response to Bacillus Calmette-Guérin (BCG) remains poorly defined. Using multispectral imaging, we characterized autonomic and sensory nerve populations within the tumor-adjacent stroma of 260 treatment-naive high-risk NMIBC tumor regions from 151 patients. Stromal tumor-associated nerve (STaN) abundance varied substantially across patients and was associated with aggressive clinicopathological features, BCG failure, and disease progression. Unsupervised clustering resolved distinct STaN phenotypic groups with differential clinical and spatial associations. Noradrenergic and neurochemically heterogeneous STaN populations showed the strongest associations with adverse outcomes. In contrast, a mixed population of nerves with high expression of the synaptic vesicle protein synaptophysin exhibited prominent spatial remodeling associated with BCG failure, despite limited prognostic value by abundance alone. Integration with an independent single-cell atlas nominated cancer-associated fibroblasts and mast cells as candidate cellular intermediaries of neural communication within the TME. Matched transcriptomic profiling revealed coordinated neuronal, extracellular matrix, lipid metabolism, and immune programs in STaN-high tumors, whereas STaN-low tumors were preferentially enriched for proliferative and hypoxic stress-response pathways. Leveraging these transcriptional features, we developed a transcriptomic classifier that distinguished STaN-high from STaN-low tumors in an independent test cohort (AUC = 0.87) and stratified clinical outcomes in additional patient cohorts. Together, these findings establish tumor innervation as a heterogeneous and clinically relevant feature of the NMIBC microenvironment, with neural abundance, phenotype, and spatial organization capturing distinct aspects of tumor behavior and clinical outcome.

## INTRODUCTION

The progression of solid tumors is shaped by interactions with the cellular and non-cellular components in and around a tumor, collectively referred to as the tumor microenvironment (TME). The TME includes diverse populations of immune cells and fibroblasts, vasculature, extracellular matrix (ECM), soluble signaling molecules, and other organ-specific, non-malignant cell types, including peripheral nerves. These components interact extensively with one another, and with tumor cells, to form a dynamic, interconnected ecosystem that shapes the local TME. The TME composition varies widely across patients and cancer types, and its evolution throughout disease progression is a well-established contributor to tumor behavior, therapeutic response, and clinical outcomes^1,2^.

Among microenvironmental components, the peripheral nervous system (PNS) innervates nearly all organs and tissues outside of the central nervous system, where it directly influences physiological and pathophysiological processes. Despite fundamental roles in tissue homeostasis and organ function, the PNS has only recently become a target of study in tumor biology^3–8^. Perineural invasion, in which tumor cells infiltrate nerve bundles, is a known behavior of invasive tumors^9,10^, but tumor innervation (local nerves infiltrating the TME) remains comparatively understudied. This gap stems, in part, from the fact that delicate nerve fibers are difficult to resolve by standard histopathology unless specifically labeled^11,12^, and because PNS neuron cell bodies reside in distant ganglia often far from the tumor site^13^, which limits their detection by transcriptomics. As a result, the neural landscape is largely hidden in routine pathological and transcriptomic workflows unless specifically probed for. Despite the growing body of evidence recognizing innervation in tumor-associated phenotypes^14–18^, its role in bladder cancer (BCa) remains poorly defined.

This gap is clinically impactful. BCa is the sixth most common cancer in the United States, with over 83,000 new cases diagnosed annually; ∼75% of patients present with non-muscle-invasive disease (NMIBC; stages Ta, Tis, and T1)^19^. NMIBC is characterized by substantial molecular and clinical heterogeneity, and high-risk cases (∼25% of diagnoses) are marked by frequent recurrence and progression^19–22^. Standard of care for high-risk patients is transurethral resection of the bladder tumor (TURBT) followed by intravesical Bacillus Calmette-Guérin (BCG) immunotherapy^19^. Although this regimen can delay or prevent progression in a subset of patients, up to 50% ultimately progress to muscle-invasive bladder cancer (MIBC; stages T2-T4)^20^, which is associated with high mortality^23,24^. Importantly, patients who initially present with NMIBC and later progress to MIBC have worse outcomes than those diagnosed with primary MIBC, highlighting the clinical importance of preventing disease progression^25–28^. Yet, there remains a fundamental gap in understanding what drives progression from non-muscle-invasive to muscle-invasive disease. Substantial work has focused on the tumor-intrinsic and microenvironmental determinants of BCG failure, improving our ability to predict response and identify patients who may benefit from earlier or alternative intervention^29,30^. As in other cancers, dynamic interactions within the NMIBC TME shape tumor behavior, and vice versa^31,32^. Therefore, a holistic understanding of how the local TME, including nerves, enables adaptation to therapy and disease progression is essential for improving outcomes in NMIBC.

The bladder is innervated by multiple peripheral nerve types with distinct functional roles^33,34^. Parasympathetic nerves release acetylcholine to promote detrusor contraction; sympathetic nerves release norepinephrine to regulate detrusor relaxation and vasodilation; and somatic motor innervation controls voluntary urination through acetylcholine release at the urethral rhabdosphincter. Sensory nerves distributed throughout the bladder wall monitor fullness and inflammation and can release neuropeptide transmitters including substance P (SP; *TAC1*) and calcitonin gene-related peptide (CGRP; *CALCA*). Each of these neurotransmitters and neuropeptides is implicated in tumor progression in other malignancies, though effects are highly cancer-, stage-, and therapy-dependent^14,15^. Notably, recent work by Lu et al. showed that sympathetic innervation drives responses to anti-PD-1 immunotherapy in MIBC^35^, raising the question of whether neural influence on immunotherapy response extends earlier in the disease course, namely to BCG treatment in the NMIBC setting. A comprehensive understanding of the diverse neural populations within the bladder TME, and their relationship to BCG response, is yet to be established.

Here, we present a comprehensive characterization of the neural microenvironment in 141 human NMIBC tumors using multispectral spatial imaging and patient-matched transcriptomic profiling to determine: ***i***) how the composition and spatial organization of sensory, sympathetic, and parasympathetic nerves within the TME is associated with BCG response and tumor biology; ***ii***) whether innervated tumors are characterized by broader changes in tumor and microenvironmental biology; and ***iii***) whether patient-matched gene expression can be leveraged to develop a transcriptomic proxy for TME innervation and improve patient risk stratification. We identify stromal innervation as a prognostic feature associated with clinical outcomes following BCG treatment and show that specific stromal tumor-associated nerve (STaN) phenotypes, particularly noradrenergic (sympathetic) and an indeterminate STaN population, are associated with poor response. Further spatial analyses reveal enrichment of noradrenergic and synaptophysin-high (SYP-high) STaNs in tumor-adjacent stromal regions consistent with putative fibroblast-rich anatomical niches. Tumors with densely innervated (STaN-high) TMEs exhibit transcriptional programs distinct from sparsely innervated tumors (STaN-low), with STaN-high tumors enriched for lipid metabolism, inflammatory, neurodevelopmental, and ECM remodeling pathways, while STaN-low tumors are characterized by proliferative, hypoxic, and stress-response signaling. To our knowledge, this study provides the most comprehensive characterization of tumor innervation in NMIBC with matched transcriptomic profiling to date.

## RESULTS

### The landscape of stromal tumor-associated nerves

A newly developed 9-color multispectral imaging panel (**Table 1**) was designed to resolve neurochemically distinct peripheral nerve populations alongside tumor and vascular architecture within single NMIBC sections (**Figure 1A**). Pan-cytokeratin (panCK) separated tumor and urothelial compartments from surrounding stroma^40^, and CD31 delineated the vasculature ^41^, establishing the spatial framework for assessing neurovascular relationships. PGP9.5 (UCHL1) and synaptophysin (SYP) identified neuronal processes ^12^ and synaptic vesicle-containing structures^42^, while tyrosine hydroxylase (TH) and vesicular acetylcholine transporter (VAChT; SLC18A3) discriminated noradrenergic (predominantly sympathetic)^43^ from cholinergic (predominantly parasympathetic) fibers^43^.

**Figure 1.**
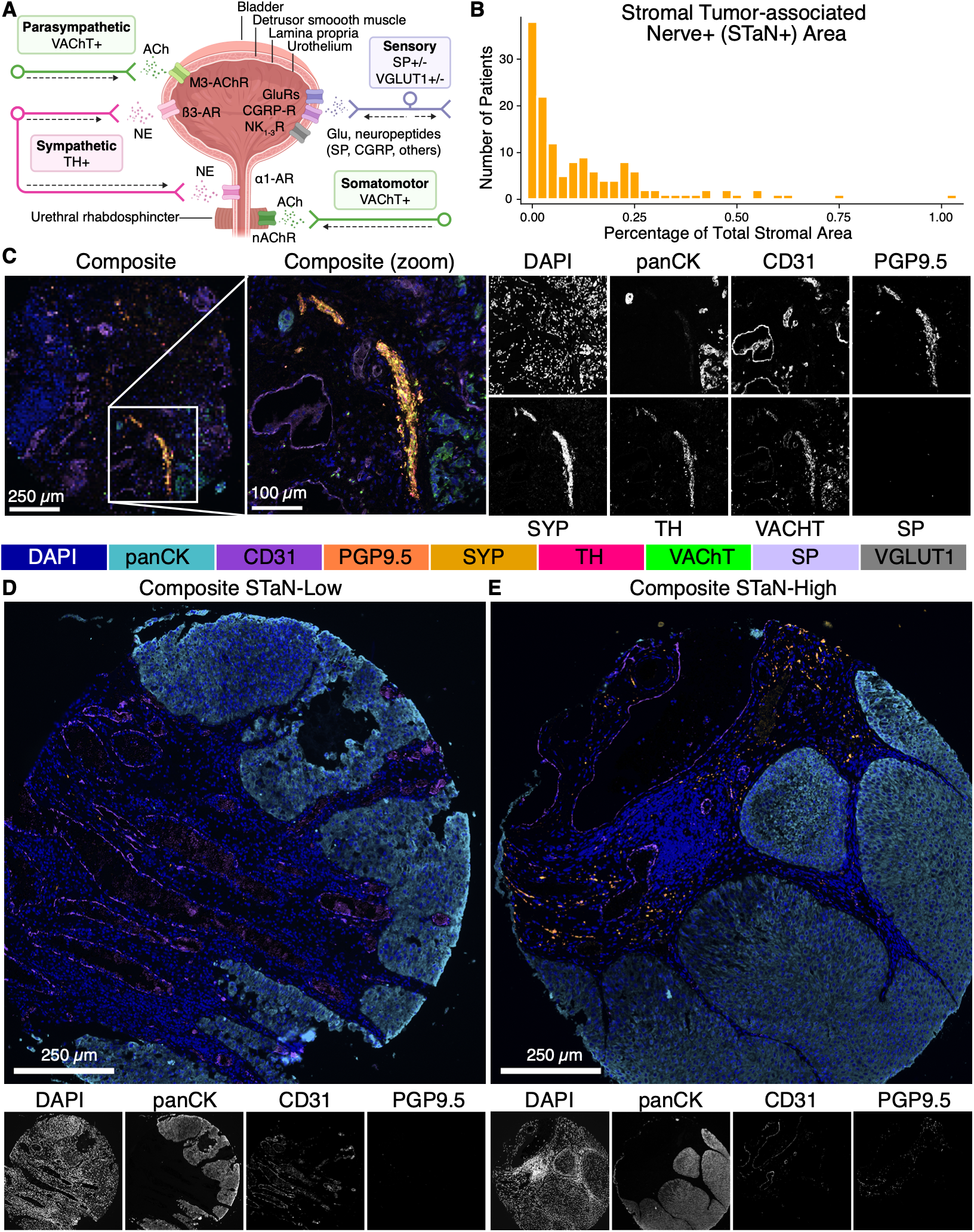
Multispectral imaging to characterize the heterogeneous neural microenvironment of human NMIBC tumors. **(A)** Simplified schematic of bladder neuroanatomy. **(B)** Histogram showing the distribution of stromal tumor-adjacent nerve (STaN+) area across the entire imaging cohort (n=151). **(C-E)** Validation of the antibody panel for STaN phenotyping. **(C)** Representative merged and single-channel images of a tumor-associated mixed nerve bundle. The composite includes PGP9.5, SYP, TH, VAChT, SP, panCK, CD31, and DAPI. **(D and E)** Representative composite images of tumors with **(D)** sparsely innervated and **(E)** densely innervated stroma. These composites include PGP9.5, SYP, panCK, CD31, and DAPI. For visualization, the display intensity for each channel was adjusted uniformly for both panels. Nerve fibers are stained with PGP9.5 (orange) and/or SYP (gold). Functional PNS subdivisions (shown only in **(C)**) are indicated by TH+ noradrenergic (hot pink), VACHT+ cholinergic (green), or SP+ sensory (lavender) nerve fibers. Tissue context is provided by panCK+ tumor cells (teal), CD31+ endothelium (dark purple), and DAPI+ nuclei (dark blue). Scale bars represent 100 µm in **(C)** and 250 µm in **(D,E)**. Images were stitched during inForm processing.

**Table 1:** 9-color nerve panel.

| Marker | Description | Gene | Function | Target |
| --- | --- | --- | --- | --- |
| panCK | Pan-cytokeratin; Recognizes a broad spectrum of epithelial cytokeratins | Several | Epithelial lineage marker | Healthy and malignant urothelial cells |
| CD31 | Cluster of differentiation 31 | PECAM1 | Endothelial cell adhesion molecule | Endothelial cells |
| PGP9.5 | Protein-gene product 9.5 | UCHL1 | Neuronal lineage marker | Pan-neuronal nerve fibers |
| SYP | Synaptophysin | SYP | Synaptic vesicle glycoprotein involved in recycling of synaptic vesicles | Synaptic nerve terminals |
| TH | Tyrosine hydroxylase | TH | Rate-limiting enzyme in catecholamine synthesis | Adrenergic (sympathetic) nerve fibers |
| VACHT | Vesicular acetylcholine transporter | SLC18A3 | Packages acetylcholine into synaptic vesicles | Cholinergic (parasympathetic) nerve fibers |
| VGLUT1 | Vesicular glutamate transporter 1 | SLC17A7 | Packages glutamate into synaptic vesicles | Myelinated mechanosensory (Aδ) nerve fibers |
| SP | Substance P | TAC1 | Neuropeptide involved in nociception and neurogenic inflammation | Many peptidergic sensory nerve fibers (Aδ and C) |
| DAPI | 4',6-Diamidino-2-phenylindole | - | Nuclear counterstain | All nuclei |

Sensory fibers, which are neurochemically diverse^36,37^ and can drive local neurogenic inflammation by releasing glutamate and nociceptive neuropeptides antidromically into the tissues they innervate^38,39^, required additional markers. CGRP and TRPV1, canonical markers of peptidergic nociceptive C fibers^37,40–42^, both gave insufficient signal-to-noise for reliable detection in this panel. We therefore used substance P (SP) to capture a subset of peptidergic sensory fibers, supported by reports of elevated circulating SP and increased tumor expression of the SP receptor TACR1 in BCa⁵¹, and VGLUT1 (SLC17A7) to distinguish myelinated Aδ fibers from unmyelinated C fibers^43^.

We applied the optimized panel to a final cohort of 260 TMA cores (0.75-1.25 mm^2^ each) from 151 patients with high-risk, treatment-naïve NMIBC who underwent TURBT (**Table 2**). The median age at diagnosis was 70 years (range, 37-88). At least 11.2% of patients were female (53% male; 35.8% unannotated), consistent with the known sex disparity in bladder cancer incidence^44^. Tumors were predominantly high-grade T1 (n=142) with a small subset of high-grade Ta (n=9), and 24 cases had concomitant carcinoma *in situ*. Of the 151 patients in the final cohort, 141 received adequate BCG therapy. Clinical annotation included BCG response (n=101 responders, n=50 non-responders), progression to muscle-invasive disease (n=29 progressors, n=122 non-progressors), longitudinal survival (n=97), and histopathological and transcriptomic subtype classification (n=97).

**Table 2:** Cohort demographics. STaN = stromal tumor-associated nerves; BCG = Bacillus Calmette-Guérin; IQR = Interquartile range; NA = not applicable; CIS = carcinoma *in situ*; LVI = lymphovascular invasion; EAU = European Association of Urology; re-TURBT = repeat transurethral resection of bladder tumor prior to BCG induction. ^a^Percentages may not add up to 100% due to rounding. ^b^p-values were calculated using complete cases only. ^c^BCG failure was defined as development of high-grade and/or muscle-invasive recurrences or persistent high-grade disease after adequate BCG therapy, consistent with international guidelines^19^.

| Characteristic<sup>a</sup> |  | All | STaN-low<br>(0-0.022% STaN+ area) | STaN-medium<br>(0.023-0.137% STaN+ area) | STaN-high<br>(0.138-1.01% STaN+ area) | p-value<sup>b</sup> |
| --- | --- | --- | --- | --- | --- | --- |
|  |  | n (%) | n (%) | n (%) | n (%) |  |
| Sex | Female | 17 (11) | 9 (6) | 6 (4) | 2 (1) | 0.039 |
|  | Male | 80 (53) | 20 (13) | 30 (20) | 30 (20) |  |
|  | Unknown | 54 (36) | 22 (15) | 14 (9) | 18 (12) |  |
| Age | Median (IQR) | 70 years<br>(63.0-77.0) | 70 years<br>(63.5-77.0) | 70 years<br>(61.5-77.0) | 72 years<br>(62.2-76.0) | 0.938 |
| Smoking | No/Stopped | 35 (23) | 13 (9) | 14 (9) | 8 (5) | 0.279 |
|  | Yes/Current | 56 (37) | 14 (9) | 21 (14) | 21 (14) |  |
|  | Unknown | 60 (40) | 24 (16) | 15 (10) | 21 (14) |  |
| re-TURBT | No | 47 (31) | 10 (7) | 18 (12) | 19 (13) | 0.09 |
|  | Yes | 104 (69) | 41 (27) | 32 (21) | 31 (21) |  |
| Adequate BCG | No | 10 (7) | 4 (3) | 4 (3) | 2 (1) | 0.772 |
|  | Yes | 141 (93) | 47 (31) | 46 (30) | 48 (32) |  |
| Stage/grade | Ta high-grade | 9 (6) | 3 (2) | 2 (1) | 4 (3) | 0.77 |
|  | T1 high-grade | 142 (94) | 48 (32) | 48 (32) | 46 (30) |  |
| T1 substage | T1 microinvasive | 21 (14) | 8 (5) | 9 (6) | 4 (3) | 0.294 |
|  | T1 extensively invasive | 73 (48) | 20 (13) | 26 (17) | 27 (18) |  |
|  | Unknown or NA | 57 (38) | 23 (15) | 15 (10) | 19 (13) |  |
| Tumor size | Small (<3cm) | 28 (19) | 10 (7) | 7 (5) | 11 (7) | 0.158 |
|  | Large (>=3cm) | 21 (14) | 11 (7) | 7 (5) | 3 (2) |  |
|  | Unknown | 102 (68) | 30 (20) | 36 (24) | 36 (24) |  |
| Tumor focality | Unifocal | 71 (47) | 29 (19) | 26 (17) | 16 (11) | 0.02 |
|  | Multifocal | 77 (51) | 21 (14) | 22 (15) | 34 (23) |  |
|  | Unknown | 3 (2) | 1 (1) | 2 (1) | 0 (0) |  |
| Concomitant CIS | Absent | 127 (84) | 50 (33) | 43 (28) | 34 (23) | < 0.001 |
|  | Present | 24 (16) | 1 (1) | 7 (5) | 16 (11) |  |
| LVI | No | 89 (59) | 28 (19) | 33 (22) | 28 (19) | 0.314 |
|  | Yes | 5 (3) | 0 (0) | 2 (1) | 3 (2) |  |
|  | Unknown | 57 (38) | 23 (15) | 15 (10) | 19 (13) |  |
| EAU risk group | High risk | 61 (40) | 25 (17) | 24 (16) | 12 (8) | 0.015 |
|  | Very high risk | 90 (60) | 26 (17) | 26 (17) | 38 (25) |  |
| BCG failure<sup>c</sup> | No | 101 (67) | 40 (26) | 36 (24) | 25 (17) | 0.006 |
|  | Yes | 50 (33) | 11 (7) | 14 (9) | 25 (17) |  |
| Progression | No progression | 122 (81) | 48 (32) | 40 (26) | 34 (23) | 0.004 |
|  | Progression | 29 (19) | 3 (2) | 10 (7) | 16 (11) |  |
| RNA-sequencing | BRS Cohort A | 68 (45) | 18 (12) | 28 (19) | 22 (15) | 0.381 |
|  | BRS Cohort B | 29 (19) | 11 (7) | 8 (5) | 10 (7) |  |
|  | Not generated | 54 (36) | 22 (15) | 14 (9) | 18 (12) |  |

Nerve fibers were localized almost entirely to the stromal compartment, as expected for NMIBC, where tumors grow along the urothelial surface overlying the innervated lamina propria. We therefore quantified stromal tumor-adjacent nerves (STaNs) as PGP9.5+ area relative to total stromal area across all cores per patient, which controls for variability in tumor-stroma composition. STaN abundance varied widely across the cohort; approximately 25% of tumors had virtually no detectable innervation (<0.01% STaN+ area), 35% were sparsely innervated (0.01-0.1%), 16% were moderately innervated (0.11-0.2%), and the remaining 24% were highly innervated (0.21-1.01%) (**Figure 1B**). Single-stain images confirmed that functionally distinct fiber types were resolvable, including within a large mixed autonomic-sensory nerve bundle (**Figure 1C**). We observed substantial STaN heterogeneity in the TME. Adjacent stroma of many tumors appeared nearly devoid of nerves (**Figure 1D**), while others were densely innervated (**Figure 1E**).

### Stromal innervation is associated with poor clinical outcomes

To determine if STaNs are associated with clinically relevant features of NMIBC, we compared STaN abundance across established clinicopathological characteristics and clinical outcomes (**Figures 2A-2G, S1, Table S1**). STaNs were not significantly associated with patient age, tumor stage (Ta vs T1), tumor location, variant histology, or FGFR3 mutation status (**Figures S1A, S1F-S1J**). Interestingly, tumors from female patients exhibited significantly lower STaN abundance than tumors from male patients (Wilcoxon, p=0.0074; **Figure S1B**). This observation raises the possibility of sex-associated differences in NMIBC innervation and warrants investigation in larger cohorts.

**Figure 2.**
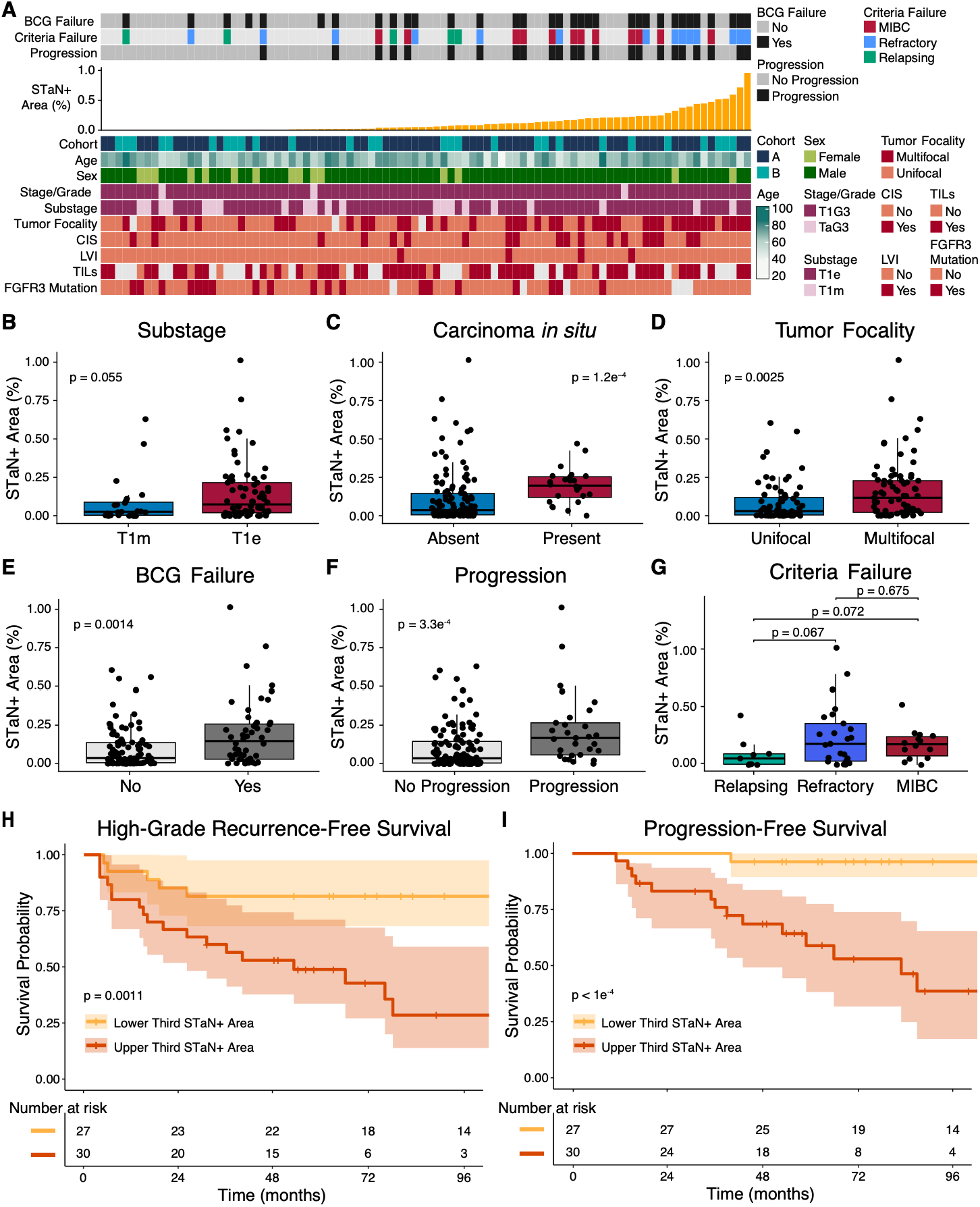
Increased STaN abundance is associated with aggressive clinicopathological features and poor clinical outcomes. **(A)** Patient-level overview of STaN abundance across patients with comprehensive clinical annotations (n=90), ordered by STaN abundance. Top annotations indicate clinical outcomes including BCG failure, failure criteria, and progression. Bottom annotations show BRS study (de Jong et al.^45^) RNA-sequencing cohort, sex, age, smoking status, stage/grade, substage, tumor size, tumor focality, presence of concomitant carcinoma *in situ* (CIS), lymphovascular invasion (LVI), tumor-infiltrating lymphocytes (TILs), and FGFR3 mutation status. **(B-G)** Associations between STaN+ area and **(B-D)** clinicopathological tumor features or **(E-G)** clinical outcomes. Aggressive clinicopathological features include **(B)** T1 substage, **(C)** concomitant carcinoma *in situ* (CIS), and **(D)** tumor focality. Clinical outcomes include **(E)** BCG failure status, **(F)** progression status, and **(G)** BCG failure criteria. Each point represents 1 patient. Boxes represent the interquartile range (IQR), center lines indicate the median, and whiskers extend to 1.5x IQR. Statistical significance was assessed using two-sided Wilcoxon rank-sum tests unless otherwise indicated. Comparisons involving more than two groups were assessed using the Kruskal-Wallis test followed by pairwise Wilcoxon rank-sum tests with Benjamini-Hochberg (BH) adjustment for multiple comparisons. P values are indicated. **(H, I)** Kaplan-Meier curves of high-grade recurrence-free survival (HG-RFS) **(H)** and progression-free survival (PFS) **(I)** stratified into upper and lower tertile STaN+ area (upper tertile, dark orange; lower tertile, gold). Shaded areas represent 95% confidence intervals. Survival differences were assessed using the log-rank test (HG-RFS p=0.0011, PFS p=6e-05). Clinical outcome and survival comparisons were restricted to patients who received adequate BCG therapy.

STaN abundance showed a consistent pattern of enrichment across clinicopathological features linked to aggressive tumor biology, including concomitant carcinoma *in situ* (CIS; Wilcoxon, p=1.2e-04; **Figure 2C**) and multifocal disease (Wilcoxon, p=0.0025; **Figure 2D**). Although incomplete clinical annotation reduced sample sizes and statistical power for several variables, similar directional trends were observed for extensive T1 invasion (T1e; Wilcoxon, p=0.055; **Figure 2B**) and prior smoking history (Wilcoxon, p=0.072; **Figure S1C**). Together, these associations link increased STaN abundance with clinicopathological features of aggressive disease.

In concordance with major urology guidelines, BCG failure was defined as persistent or recurrent high-grade disease after 6 months of adequate BCG induction^19^. Consistent with aggressive disease features, STaN abundance was significantly higher in patients who later failed to respond to BCG treatment (Wilcoxon, p=5.3e-04, **Figure 2E**) and those who progressed to muscle-invasive disease (Wilcoxon, p=2.1e-04; **Figure 2F**).Within cases of BCG failure, STaN abundance at diagnosis was significantly higher in tumors from patients with BCG-refractory disease (persistent or recurrent high-grade disease during BCG treatment) than in those with BCG-relapsing disease (high-grade recurrence following an initial response to BCG; Wilcoxon, p=0.032; **Figure 2G**), suggesting that STaN abundance may distinguish different patterns of BCG treatment failure.

To evaluate the clinical significance of STaN-high and STaN-low TMEs in NMIBC, we compared longitudinal survival outcomes between patients in the upper and lower STaN tertiles defined across patients who received adequate BCG therapy in the final imaging cohort (n=141 out of n=151). High-grade recurrence-free survival (HG-RFS) and progression-free survival (PFS) were significantly worse in patients with STaN-high tumors than those with STaN-low tumors (HG-RFS log-rank, p=0.0011; PFS log-rank, p=6e-05; **Figures 2H, 2I**). Survival patterns across all three tertiles and using median-based stratification were consistent with the primary analysis (**Figure S2**). Together, these findings support STaNs as a characteristic of aggressive tumors and a prognostic feature of high-risk NMIBC.

### High nerve abundance correlates with multiple high-risk molecular subtypes

Because STaN abundance was associated with aggressive clinicopathological features and adverse clinical outcomes, we next asked whether this information was redundant with existing clinicopathological^23,46^ and transcriptomic^45,47–49^ classification systems, or varied independently of them. Accordingly, we compared STaN abundance across multiple established NMIBC classification frameworks (**Figures 3A, S3; Table S1**). STaNs were enriched within the highest-risk EAU^23^ and EORTC^46^ clinicopathological risk groups, consistent with their association with aggressive disease. In contrast, STaNs were not preferentially enriched in any established transcriptomic molecular subtype (**Figures 3B, S3**), indicating that STaNs are not simply a surrogate for existing transcriptional classifications. Importantly, STaN abundance provided additional prognostic resolution across both clinicopathological risk groups (EAU and EORTC; **Figures 3E, 3F, S3**) and within multiple high-risk molecular subtypes, including BRS3^45^, UROMOL Class 2a^49^, and the Chicago T1 NMIBC luminal genomically unstable (LumGU; PFS only) and MYC-like subtypes^50^ (**Figures 3C, 3D**). Within these high-risk groups, patients with STaN-high tumors consistently had worse clinical outcomes than those with STaN-low tumors. In contrast, little or no additional stratification was observed within several lower-risk subtype groups. These findings demonstrate that STaNs capture a biological axis not captured by existing molecular subtype classifications while providing complementary prognostic information that further refines risk stratification in high-risk NMIBC.

**Figure 3.**
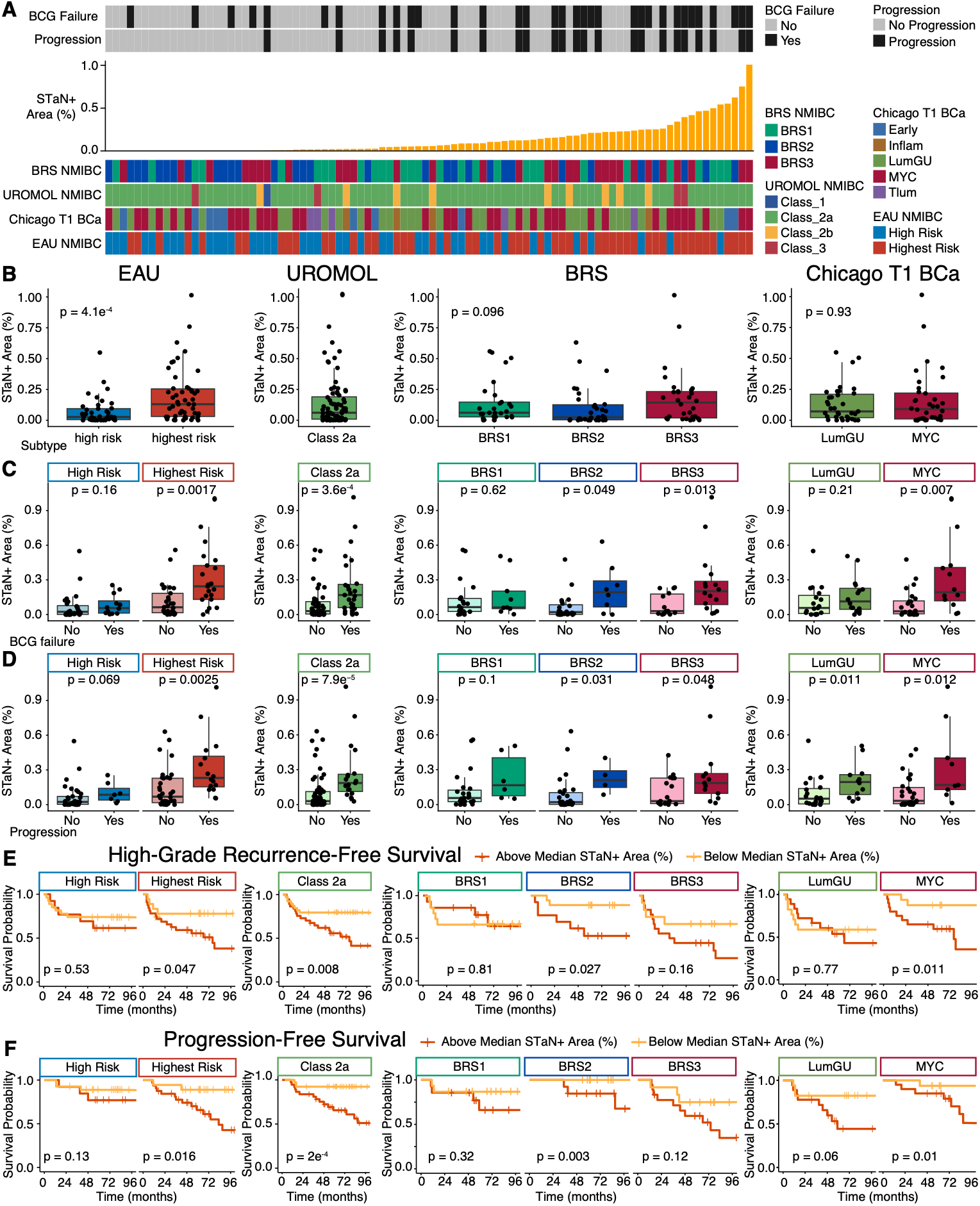
STaN abundance further stratifies clinical outcomes in established histopathological molecular subtypes. **(A)** Patient-level overview of STaN+ area alongside clinical outcomes and multiple clinicopathological (EAU) and molecular (UROMOL, BRS, Chicago T1) NMIBC classification systems. Patients are ordered by STaN+ area (%). Clinical endpoints (BCG failure/high-grade recurrence, and progression) and subtype assignments are shown as annotated tracks. **(B)** STaN abundance according to subtype classifications with >=10 patients total and >=3 patients per clinical outcome. **(C,D)** STaN+ area according to **(C)** BCG failure status and **(D)** progression status within subtypes with >= 10 patients total and >=3 patients per outcome. Boxes indicate the interquartile range (IQR), center lines denote the median, and whiskers extend to 1.5x IQR. Each data point represents a single patient. Statistical significance was assessed using the Wilcoxon rank-sum test or Kruskal-Wallis test for groups with more than two comparisons. P-values are indicated. **(E-F)** Kaplan-Meier survival analyses of **(E)** high-grade recurrence-free survival (BCG failure; HG-RFS) and **(F)** progression-free survival (PFS) stratified by median STaN+ area within each subtype. Median stratification was used in place of upper/lower tertiles to increase sample sizes per group. Statistical significance was assessed using the log-rank test. P-values are indicated. Clinical outcome and survival comparisons were restricted to patients who received adequate BCG therapy.

### STaN-high and STaN-low tumors exhibit distinct microenvironmental and transcriptional phenotypes

To define the transcriptional programs associated with innervated TMEs, we performed differential gene expression analysis comparing STaN-high and STaN-low tumors with matched transcriptomic data (**Figure 4; Tables S2**). STaN-high tumors preferentially expressed genes involved in extracellular matrix remodeling, inflammation, neuronal adhesion, axon guidance, and neurodevelopment, including *NRCAM*, *NRXN1*, *DNER*, *C1QL1*, *PTPRT*, *GAS1*, and *REEP1*, together with developmental regulators such as *PAX6*, *PRDM16*, and *FOXN4* (**Figure 4A; Table S2**). Consistent with these transcriptional changes, functional gene set enrichment analysis identified coordinated enrichment of neuronal signaling, synaptic organization, lipid metabolism, and immune-associated pathways in STaN-high tumors (**Figures 4B, 4C, S4A; Table S2**). Hallmark gene set analysis revealed a distinct pattern in STaN-low tumors, with enrichment of proliferative and cellular stress response programs, including MYC targets, MTORC1 signaling, hypoxia, and the unfolded protein response (**Figure 4C**). Cell type signature enrichment analysis further demonstrated enrichment of diverse immune, stromal, vascular, and neural cell-associated transcriptional signatures in STaN-high tumors, suggesting a more compositionally heterogeneous and complex microenvironment (**Figure S4B**). These coordinated transcriptional profiles suggest that differences in STaN abundance reflect distinct biological states characterized by broader transcriptional and compositional remodeling of the TME, rather than isolated differences in innervation.

**Figure 4.**
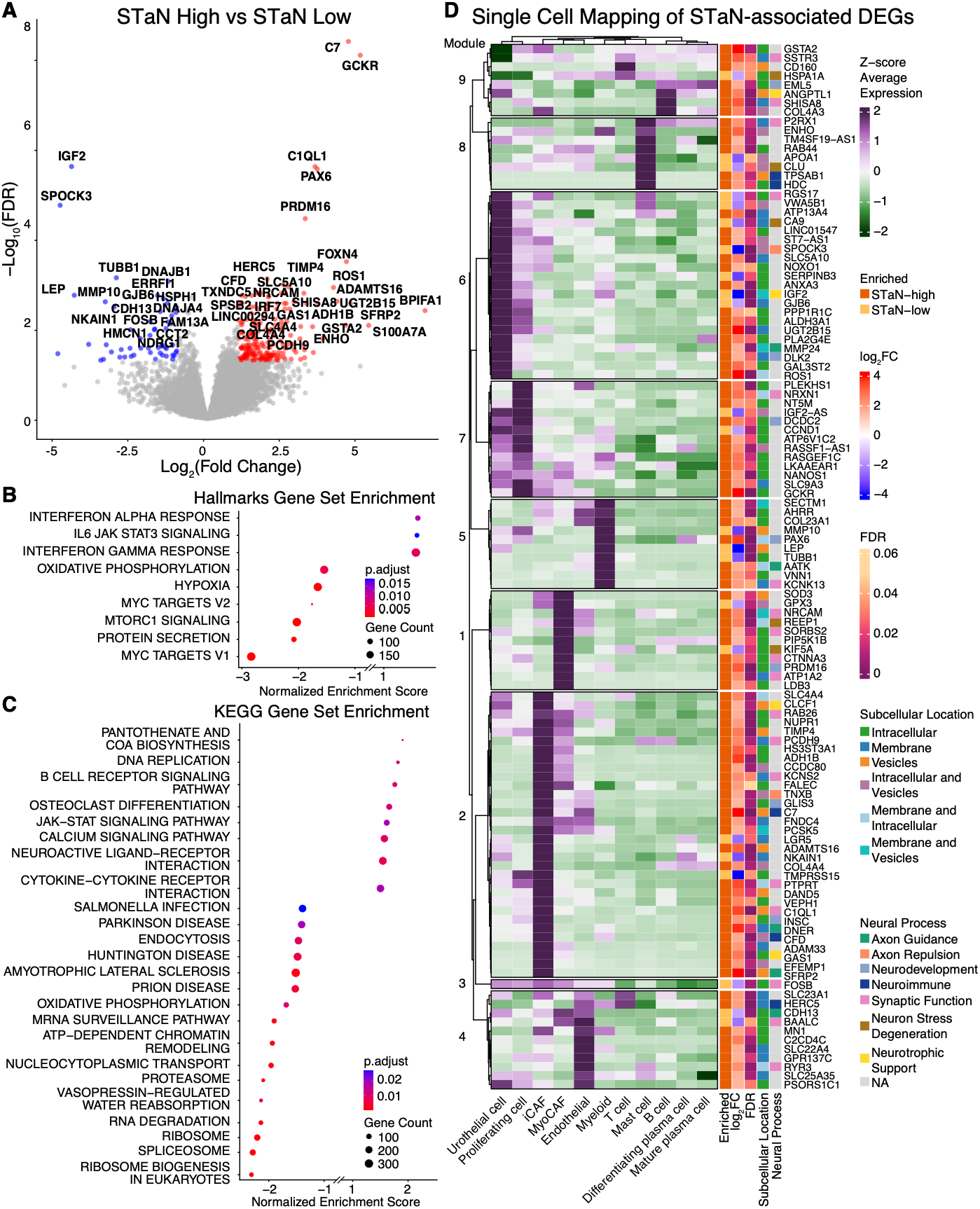
STaN-high tumors exhibit distinct transcriptional states. **(A)** Volcano plot of differentially expressed genes (DEGs) between tumors in the upper (STaN-high; n=33) and lower (STaN-low; n=33) STaN tertiles. STaN tertiles were defined within each sequencing cohort (A or B) from the BRS study (de Jong et al.^45^). Differential expression analysis was performed using DESeq2^51^ with sex and cohort included as covariates. Significantly upregulated (red; n=220) and downregulated (blue; n=53) genes in STaN-high tumors are shown (FDR <0.05, |log2FC| >1). A total of 17,565 protein-coding, microRNA, and long non-coding RNA genes were evaluated. **(B, C)** Gene set enrichment analysis (GSEA) of **(B)** MSigDB Hallmark and **(C)** KEGG pathways using the DESeq2 Wald statistic to generate a pre-ranked gene list. Dot size represents gene set size, and color denotes normalized enrichment score (NES). Positive NES indicates enrichment in STaN-high tumors, whereas negative NES indicates enrichment in STaN-low tumors. (**D**) Expression of significant STaN-associated DEGs across major cell populations from an independent published single-cell bladder cancer atlas (Chen et al.^52^; PRJNA662018). Rows represent DEGs identified in panel A and columns represent annotated cell populations. Heatmap color indicates scaled average expression. Gene annotations indicate STaN enrichment group (STaN-high or STaN-low), log2 fold-change (log2FC), statistical significance, predicted subcellular localization (Human Protein Atlas), and curated functional categories including extracellular matrix remodeling, neuronal development, axon guidance, cell adhesion, neurotransmission, and signaling. This analysis was performed to provide cellular context for STaN-associated transcriptional programs and was not used to validate the imaging cohort.

To provide cellular context for STaN-associated transcriptional changes, we mapped statistically significant DEGs onto an independent single-cell bladder cancer atlas (**Figure 4D, S5**). Looking gene by gene, each STaN-associated DEG is shown against the tumor and TME populations in bladder tumors, mapping each gene to its likely cell type. It does not infer direct interactions between nerves and those cells, nor does it independently validate the association between these genes and STaN abundance. The data show that DEGs do not distribute evenly across cell types. Several transcriptional programs enriched in STaN-high tumors converged on a small number of stromal and immune populations, with inflammatory cancer-associated fibroblasts (iCAFs) and mast cells accounting for much of the STaN-high enriched signal.

The iCAF-associated genes fell into three functional groups: extracellular matrix remodeling (*ADAMTS16, TIMP4, SFRP2*), developmental regulation (*PRDM16, GAS1*), and neuronal adhesion and axon guidance (*NRCAM, DNER, PTPRT, SEMA5B*), which is of interest given the neural phenotype in this study. Mast cells showed a different and more directional pattern. Genes upregulated in STaN-high tumors mapped to canonical effector genes, including the tryptase *TPSAB1*, the histamine-synthesizing enzyme *HDC*, and the degranulation-associated GTPase *RAB44*, suggesting enhanced effector and secretory capacity. The mast-cell-expressed genes associated with STaN-low tumors were different in character: *CLU, RGS17*, and *HSPA1A* are regulatory and stress-response genes expressed broadly across cell types, rather than markers of effector function. Both groups therefore contain mast cell genes, but they represent different mast cell programs.

Together, these findings indicate that STaN-associated transcriptional differences span multiple cellular compartments of the TME, involving stromal and immune populations in addition to tumor cells, and reflect broader microenvironmental biology beyond canonical peripheral nerve signaling. These findings identify a robust transcriptional phenotype associated with STaN abundance, motivating the development of a transcriptome-based classifier to predict TME innervation.

### A 12-gene signature predicts STaN-high and STaN-low tumors across independent cohorts

There are numerous computational methods to estimate the abundance of tumor, stromal, fibroblast, and immune cell populations from bulk transcriptomic data^53^. However, existing approaches to infer neural content rely on predefined or computationally derived neural gene signatures, and it remains unclear to what extent these signatures reflect actual tumor innervation^54–59^. We therefore developed a transcriptomic classifier to distinguish STaN-high and STaN-low tumors from bulk gene expression using patient-matched quantitative histologic measurements of STaN abundance as image-derived ground truth (**Figures 5, S6-S8**).

**Figure 5.**
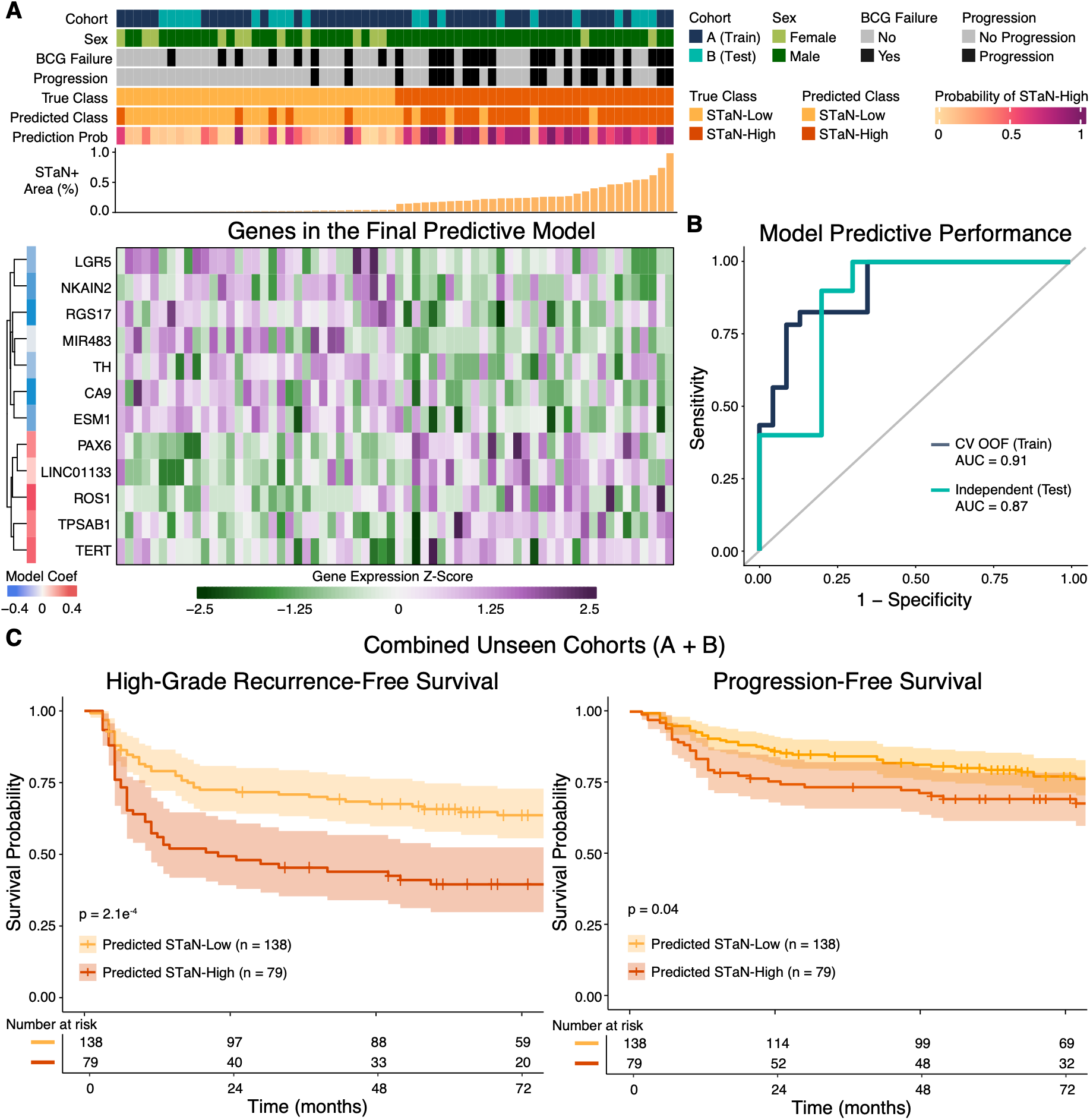
Bulk transcriptomic profiles accurately predict STaN abundance. **(A)** Heatmap showing expression of genes with nonzero coefficients in the final elastic net model across two independent RNA-sequencing cohorts from the BRS study (de Jong et al.^45^), which were sequenced on different platforms. BRS cohort A was used for model training and BRS cohort B for independent evaluation. Patients (columns) are ordered by imaging-derived STaN abundance. Gene expression data from patients with matched bulk RNA sequencing and imaging were used to train and evaluate a classifier predicting STaN-high (upper tertile) versus STaN-low (lower tertile) tumors, with STaN-positive stromal area (%) serving as the ground truth. To increase sample size and balance the classes, STaN tertiles were calculated separately within each cohort (training: n = 23 STaN-low and n = 23 STaN-high; test: n = 10 STaN-low and n = 10 STaN-high). **(B)** Receiver operating characteristic (ROC) curves for cross-validated out-of-fold predictions in the training cohort (AUC = 0.909, sensitivity = 0.826; specificity = 0.870; accuracy = 0.848; n = 46) and predictions in the independent test cohort (AUC = 0.870; sensitivity = 0.800; specificity = 0.800; accuracy = 0.800; n = 20). **(C,D)** Clinical relevance of model predictions. Kaplan-Meier analyses of **(C)** high-grade recurrence-free survival (HG-RFS) and **(D)** progression-free survival (PFS) were performed in a pooled cohort of patients not used for model development. This unseen cohort (n=217) included patients classified as STaN-medium by imaging and therefore excluded from training and testing (middle tertile; n = 31), as well as patients without matched imaging-based STaN quantification (n=186). Patients were stratified by their model-predicted STaN class (n=79 STaN-high; n=138 STaN-low), and survival differences were assessed using the log-rank test.

The classifier was developed using a subset of the two patient cohorts from the BRS study (de Jong et al.^45^) with matched STaN imaging and RNA-sequencing data (**Figure 5A, S6**). BRS Cohort A was used for model training, while BRS Cohort B, which was sequenced on a different platform, served as an independent test cohort. The final elastic net model retained 12 genes and accurately distinguished STaN-high from STaN-low tumors, achieving an out-of-fold AUC of 0.91 (accuracy = 0.848) in BRS Cohort A (train) and maintaining strong performance in BRS Cohort B (test; AUC = 0.87; accuracy = 0.800; **Figures 5A, 5B, S7, S8; Table S3**). Notably, the final 12-gene signature consisted largely of non-neuronal genes, consistent with our observation that STaN abundance is associated with coordinated remodeling of the TME rather than isolated expression of neuronal genes or neurotransmitter receptors.

To evaluate the clinical relevance of transcriptomically inferred innervation, we applied the classifier to unseen patients from the BRS cohort and independent external NMIBC cohorts lacking matched imaging-based STaN quantification. Consistent with comparisons using measured STaN abundance to stratify patients, predicted STaN-high tumors exhibited significantly worse high-grade recurrence-free survival (log-rank, p= 4.681e-05) and progression-free survival (log-rank, p=0.0146) compared to STaN-low tumors in an unseen internal cohort (**Figure 5C, 5D**). Predicted STaN status similarly stratified BCG failure in independent external cohorts (log-rank, p=0.021; **Figure S8E**). Due to the limited number of progression events, progression-free survival could not be robustly evaluated in the external cohorts. However, predicted STaN status provided additional prognostic resolution for BCG failure and progression within established clinicopathological and molecular subtypes of NMIBC (**Figure S9**). Together, these findings establish proof of principle that imaging based STaN abundance can be inferred from bulk transcriptomic data, supporting its further evaluation as a biomarker of BCG failure and disease progression.

### Distinct nerve phenotypes are prognostic for BCG failure and progression

Although patients with STaN-high tumors exhibit poor clinical outcomes, it is unclear whether this relationship reflects all peripheral nerves or specific neural populations. We therefore classified molecularly distinct STaN phenotypes within segmented PGP9.5+ nerve imaging “objects” by K-means clustering of multiplexed immunofluorescence marker expression (mean pixel intensity, MPI) (**Figures 6A, 6B; Table S4**). Clustering identified seven distinct subpopulations corresponding to canonical autonomic, sensory, and pre-synaptic phenotypes, including cholinergic/parasympathetic (cluster 4; VAChT+), noradrenergic/sympathetic (cluster 2; TH-mid and cluster 5; TH-high), and sensory (cluster 1; VGLUT1+/SP+) populations. We also detected a mixed synaptophysin-high population (SYP-high; cluster 6; SYP^+^/SP^-^/VGLUT1^+/-^ /TH^+/-^/VAChT^+/-^), consistent with a heterogeneous population of SYP-high autonomic and sensory nerve fibers (**Figures 6C, S10**). A seventh population (cluster 7; indeterminate) represented a heterogeneous population of nerve fibers lacking sufficient expression for confident phenotype assignment. Inspection of spatial reconstructions of the segmented nerve masks suggested that this population comprised both genuine nerve phenotypes not captured by the canonical autonomic and select sensory markers included in our panel, as well as segments with sub-threshold marker expression contiguous with otherwise classifiable autonomic or sensory nerves. Accordingly, clusters 6 and 7 were interpreted as heterogeneous residual populations rather than discrete neuronal subtypes (**Figures 6A, 6C**). Manual inspection of representative images revealed that cluster 3 consisted of nucleated PGP9.5+/SYP+/VGLUT1+ cells with morphology consistent with fibroblasts rather than nerve fibers; this cluster was excluded from all downstream analyses. Tumor-adjacent stroma predominantly included clusters 2 (TH-mid), 6 (SYP-high), and 7 (Indeterminate) (**Figure 6D, 6E, S10**). Notably, the STaN landscape was enriched for noradrenergic, SYP-high, and a mixed/unclassified STaN population, whereas cholinergic fibers were comparatively sparse, suggesting selective remodeling of peripheral nerve composition within the bladder TME.

**Figure 6.**
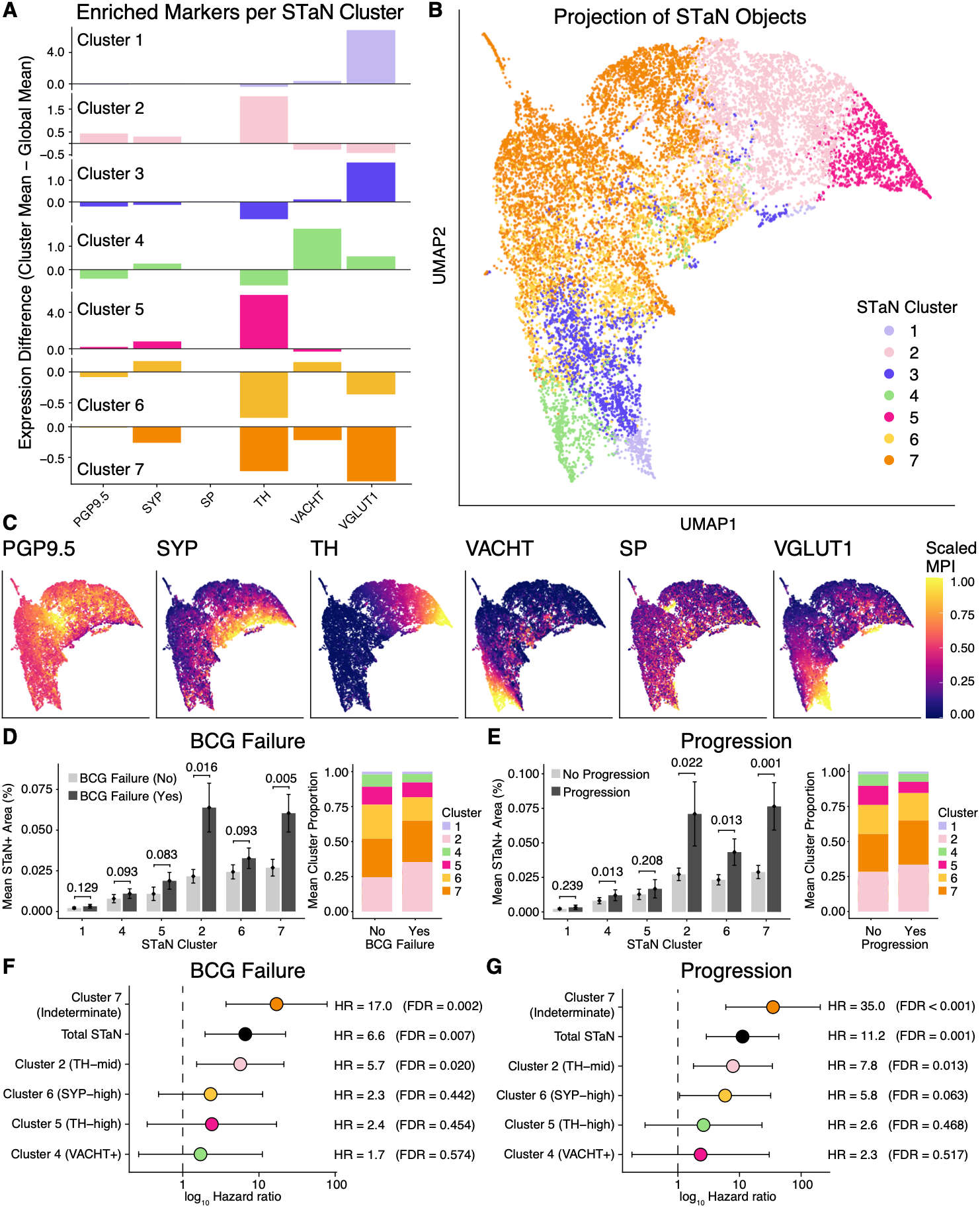
Distinct STaN phenotypes differentially associate with clinical outcome. **(A)** Cluster centroid profiles showing relative marker expression (mean pixel intensity, MPI) for each nerve cluster relative to global mean intensity. Clusters were annotated based on relative marker expression and correspond to putative sensory (VGLUT1+/SP+; cluster 1), cholinergic/parasympathetic (VAChT+; cluster 4), noradrenergic/sympathetic (TH+; clusters 2 and 5), and mixed/indeterminate populations including SYP-high (cluster 6) and marker-negative STaNs (cluster 7). **(B)** Uniform manifold approximation and projection (UMAP) of segmented PGP9.5+ STaN objects from the final imaging cohort (n=15,609 candidate STaN objects from n=260 cores and n=151 patients), colored by unsupervised K-means cluster assignment (k=7). Clustering and UMAP embedding were performed using all candidate STaN objects prior patient-level exclusions; the visualization shown here was restricted to STaN objects from the final imaging cohort. **(C)** UMAPs colored by MPI of individual nerve markers (PGP9.5, SYP, TH, VAChT, SP, VGLUT1), showing the distribution of marker expression across clusters. Marker MPI was scaled relative to all segmented objects prior to filtering and STaN clustering. Cluster assignment and marker expression for filtered and unfiltered objects are available in **Table S4**. **(D,E)** STaN cluster composition across **(D)** BCG response groups (BCG failure vs no failure; n=44 and n=97, respectively) and **(E)** progression status (progression vs no progression; n=25 and n=116, respectively) Left, mean STaN+ area (%) per cluster; right, proportional composition of STaN clusters per outcome. Bars represent mean +/-SEM. P-values were calculated using Wilcoxon rank-sum tests and adjusted using the Benjamini-Hochberg procedure. **(F,G)** Multivariable Cox proportional hazards models for **(F)** BCG failure and **(G)** progression including total STaN and STaN phenotype abundances, adjusted for age. Stage and grade were not adjusted for due to the homogenous nature of the cohort.

We next examined the relationship between individual STaN phenotypes and clinical outcomes. Noradrenergic cluster 2 (TH-mid) and the mixed/indeterminate cluster 7 were significantly associated with BCG failure and disease progression (**Figures 6D, 6E**). Clusters 2 and 7 remained associated with BCG failure and progression in Cox proportional hazards models adjusted for age (**Figures 6F, 6G**). Interestingly, STaN cluster 6 (SYP-high) was significantly enriched in disease progression (FDR=0.016) but not BCG failure (**Figures 6D-6G**). Although the molecular identity of cluster 7 could not be resolved with the current marker panel, its strong and independent associations with BCG failure and progression identify it as a clinically relevant nerve population warranting further molecular characterization.

These findings indicate that the adverse clinical associations of STaNs are not uniformly distributed across peripheral nerve phenotypes. TH-mid noradrenergic and a mixed/indeterminate population of STaNs emerged as the strongest predictors of both BCG failure and disease progression. Conversely, STaNs enriched for the canonical synaptic terminal marker SYP corresponded specifically with progression. These results suggest that individual STaN phenotypes carry different clinical associations and are associated with distinct aspects of tumor behavior, highlighting the importance of resolving neural phenotype rather than treating tumor innervation as a homogeneous feature of the TME.

### Spatial organization of STaNs suggests a multicellular model of neural–tumor communication

To determine whether STaNs preferentially localize to distinct anatomical niches, we quantified STaN phenotype-specific spatial enrichment relative to the tumor-stroma border in STaN-high tumors (**Figure 7A**). In general, SYP-high STaNs preferentially localized within tumor-adjacent stromal regions, whereas other nerve populations showed comparatively uniform spatial distributions (**Figure 7B; Table S5**). While overall STaN abundance identified prognostically relevant nerve populations, spatial analysis revealed an additional layer of STaN organization associated with BCG response. The TH-mid noradrenergic population (cluster 2) was significantly enriched near the tumor border (26-50 μm; FDR = 0.03) in tumors that subsequently failed BCG therapy, consistent with its independent association with adverse clinical outcome (**Figures 7C, 7D**). SYP-high STaNs (cluster 6) were similarly redistributed toward the tumor-adjacent stroma in BCG-failing tumors, with enrichment immediately adjacent to the tumor border (0-25 μm; FDR = 0.023) and relative underrepresentation in the distal 201-300 μm compartment (FDR = 0.00014). SYP-high STaNs were also relatively underrepresented within the distal 201-300 μm compartment in progressing tumors (FDR = 0.031), although preferential localization toward the tumor border did not reach statistical significance in more proximal bands. These spatial patterns occurred despite the absence of an association between overall SYP-high STaN abundance and BCG failure, suggesting that spatial organization captures information about BCG response not apparent from abundance. Conversely, the indeterminate population (cluster 7) exhibited no significant spatial redistribution despite being the strongest predictor of both BCG failure and progression by overall abundance (**Figures 7C, 7D**), indicating that clinically relevant features differ among STaN phenotypes, with some relationships captured primarily by abundance and others by spatial organization.

**Figure 7.**
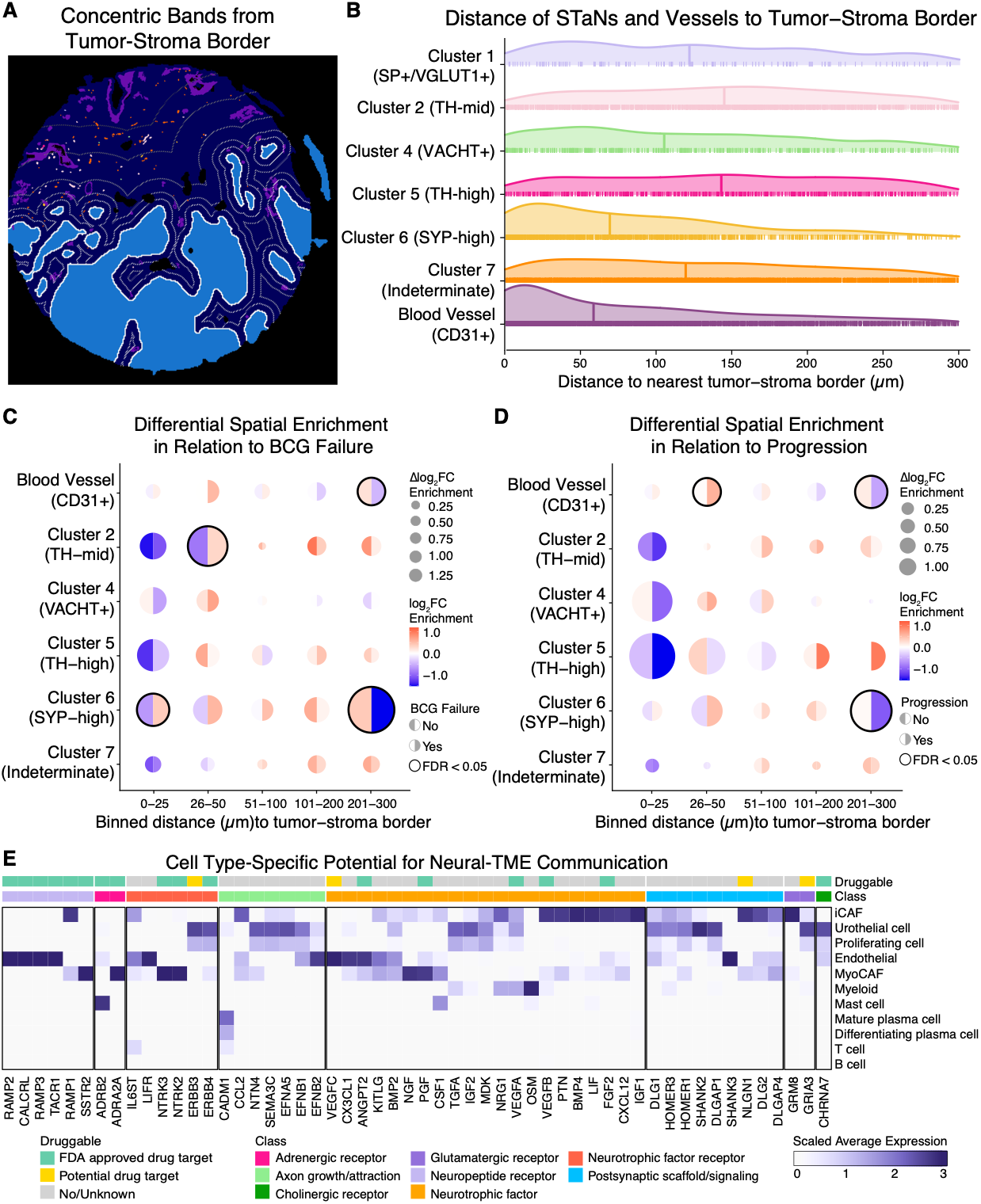
Spatial organization of STaNs reveals clinically relevant proximity patterns and nominates candidate cellular partners for neural-TME communication. **(A)** Representative tissue mask illustrating concentric stromal distance bands (0-25, 26-50, 51-100, 101-200, and 201-300 μm) generated relative to the tumor-stroma border for spatial enrichment analysis. **(B-D)** Spatial analyses of STaN and vessel objects in relation to the tumor-stroma border. **(B)** Spatial density distributions of masked vessels and individual STaN phenotypes as a function of distance from the tumor-stroma border. Curves represent the distribution of segmented nerve objects across all images in spatial cohort. **(C,D)** Differential spatial enrichment of STaN phenotypes in relation to **(C)** BCG failure and **(D)** progression. Circle size represents the magnitude of the difference in log2 fold-change (Δlog2FC) enrichment between outcome groups, while circle color indicates the direction and magnitude of enrichment (log2FC). Split circles depict enrichment estimates within each outcome group (No vs Yes), and black outlines indicate FDR-adjusted significance (FDR < 0.05). Enrichment was calculated from STaN+ area (µm^2^) relative to stromal area (µm^2^) within each distance band, with overall STaN abundance within 300 µm of the tumor-stroma border accounted for in the statistical model. Spatial analyses were restricted to STaN-high tumors with stromal tissue at least 300 μm from the tumor-stroma border (n=45 images from n=32 patients). Cluster 1 was excluded from the spatial enrichment models because of its low sample size, which led to non-identifiable (singular) mixed-effects model fits. Raw spatial metrics and modeling statistics are available in **Table S5**. **(E)** Cell type-specific expression of candidate genes for mediating bidirectional communication between STaNs and the TME mapped to an independent published single-cell RNA-sequencing atlas of two high-grade NMIBC tumors (Chen et al.^52^; PRJNA662018). Heatmap showing the normalized average expression of curated genes involved in neurotransmitter signaling, neuropeptide signaling, neurotrophic signaling, axon guidance, and synaptic communication across annotated cell populations. Genes with < 0.25 normalized average expression were excluded. Genes are grouped by functional class and annotated according to druggability. Normalized expression values were globally scaled for visualization. The atlas was used to identify candidate cell populations capable of reciprocal neural communication and was not used as validation of the imaging cohort.

Spatial remodeling of STaNs was accompanied by a similar shift in the stromal vasculature (CD31+). In the BCG failure group, vessels were less concentrated in the distal 201-300 μm compartment (FDR = 0.015). In the progression group, vessels were more concentrated near the tumor border (26-50 μm; FDR = 0.023) and less concentrated in the distal 201-300 μm compartment (FDR = 0.0074), consistent with a redistribution of the stromal vasculature toward the tumor-adjacent niche (**Figures 7C, 7D**). Together, these findings localize specific outcome-associated changes in STaN and vascular organization to the tumor-adjacent stroma rather than uniformly throughout the TME. Notably, the 0-50 μm stromal compartment corresponds anatomically to the superficial lamina propria immediately beneath the urothelium, a region enriched in resident fibroblasts in normal bladder^60^ and CAFs in BCa^61^. This observation prompted us to investigate which cell populations within the TME express molecular machinery compatible with reciprocal neural communication.

We therefore integrated an independent published single-cell RNA-sequencing atlas of bladder urothelial carcinoma (PRJNA662018)^52^. Rather than serving as validation of our imaging findings, this external dataset was used to identify cell populations expressing genes compatible with neural communication. iCAFs emerged as a prominent candidate for STaN-TME communication, exhibiting broad expression of secreted neurotrophic and growth factor ligands, including *VEGFB*, *PTN*, *BMP4*, *LIF*, *FGF2*, *CXCL12*, and *IGF1*, together with postsynaptic signaling machinery including *DLG1*, *HOMER3*, *HOMER1*, *NLGN1*, and *DLG2*. iCAFs also prominently expressed many STaN-high-associated DEGs identified in **Figure 4**, further supporting this stromal population in neural remodeling. Urothelial cells similarly expressed axon-guidance and cell-interaction molecules, including *SEMA3C*, *EFNA5*, *EFNB1*, and *EFNB2*, as well as postsynaptic scaffold proteins including *DLG1*, *HOMER3*, *HOMER1*, *SHANK2*, and *DLGAP1*. In contrast, urothelial cells showed little expression of canonical PNS neurotransmitter receptors, with the exception of the glutamate receptor *GRIA3* and cholinergic receptor *CHRM7*.

Noradrenergic receptor expression was likewise largely absent from urothelial cells and most other cell populations despite the strong prognostic associations of noradrenergic STaNs, with expression primarily restricted to myoCAFs and mast cells. MyoCAFs expressed the α2A-adrenergic receptor (*ADRA2A*) together with nerve growth factor (*NGF*), identifying a potential reciprocal signaling axis through which myoCAFs could both respond to sympathetic input and support neural recruitment or maintenance. Several of the strongest STaN-high-associated DEGs and pathways were mast cell-enriched (**Figures 4D, S4B**), and mast cells expressed the β2-adrenergic receptor (*ADRB2*), identifying another plausible cellular target of noradrenergic signaling (**Figure 7E**).

Although these scRNA-seq analyses were performed on an external cohort lacking imaging-based STaN quantification, their convergence with the bulk transcriptomic and spatial findings supports a broader neuro-immune-stromal-tumor communication network involving iCAFs, myoCAFs, mast cells, and urothelial cells. In particular, the presence of postsynaptic machinery in urothelial cells, together with the preferential localization of SYP-high STaNs immediately adjacent to the tumor border, raises the intriguing possibility that nerve-tumor communication may include short-range or contact-dependent interactions distinct from canonical diffuse neurotransmitter signaling.

These findings demonstrate that STaNs are not biologically homogeneous. Distinct STaN populations display unique molecular, spatial, and prognostic characteristics, with noradrenergic and putative synaptically active populations contributing through phenotype-specific patterns of abundance and spatial organization. Collectively, our findings suggest that STaN abundance should be viewed not simply as a measure of microenvironmental innervation, but as a systems-level feature reflecting coordinated remodeling of the neural, stromal, vascular, and transcriptional microenvironment.

## DISCUSSION

Despite growing recognition of neural remodeling within the TME, the composition, organization, and clinical significance of tumor innervation in NMIBC remain poorly understood. Here, we integrated multiplex spatial imaging, transcriptomics, computational pathology, and machine learning to define the STaN landscape in human NMIBC and investigate its relationship with tumor biology and clinical outcomes. STaN abundance was highly heterogeneous and stratified risk of BCG failure and disease progression, while STaN-high tumors showed distinct transcriptional programs, selective remodeling of STaN phenotype composition, and spatial organization. Together, these findings suggest that STaN abundance reflects a coordinated microenvironmental state, not simply increased tumor innervation, providing a systems-level framework for understanding neural contributions to therapeutic response and disease progression in NMIBC.

The adverse clinical associations of STaNs were not explained equally by all peripheral nerve populations. The strongest prognostic signal arose from an indeterminate population (cluster 7), followed by the noradrenergic TH-mid population (cluster 2), both of which were independently predicted BCG failure and progression. While some phenotypes were primarily informative through overall abundance, others, like the mixed population of SYP-high STaNs (cluster 6), derived much of their clinical relevance to BCG failure from spatial organization. In contrast, overall SYP-high STaN abundance was more closely associated with disease progression. Together, these findings demonstrate that neural identity, abundance, and spatial organization each provide complementary information regarding the biological role of tumor innervation in NMIBC.

The predominance of noradrenergic and a mixed SYP-high nerve population within tumor-adjacent stroma also contrasts with the normal bladder submucosa, which is reported to be innervated primarily by sensory fibers^33,34,62^. Although healthy bladder tissue was not examined in the present study, this pattern implies selective remodeling of STaN composition during tumor development rather than generalized expansion of all neural populations.

The preferential association of the TH-mid, but not TH-high, noradrenergic population with adverse clinical outcomes is notable given the established role of noradrenergic signaling in regulating tumor progression and anti-tumor immunity. Although the biological basis for this distinction remains unclear, it suggests that functionally distinct states of sympathetic innervation exist within the TME. One possibility is that TH-mid fibers represent actively remodeling or tumor-responsive sympathetic axons, whereas TH-high fibers reflect more mature, established sympathetic innervation^63^. Consistent with this hypothesis, TH-mid fibers showed stronger spatial enrichment near the tumor border and more robust associations with adverse clinical outcomes. These processes provide plausible routes by which localized sympathetic remodeling could shape BCG response.

STaN-high tumors differed from STaN-low tumors far beyond their neural content alone. Our transcriptional analyses suggest that STaN abundance reflects fundamentally distinct biological states and coordinated TME remodeling involving multiple stromal and immune compartments. The enrichment of neurodevelopmental, extracellular matrix, and immune pathways in STaN-high tumors, together with the scarcity of canonical neuronal genes in the transcriptomic classifier, argues that the biological consequences of tumor innervation extend beyond direct nerve-tumor communication to involve broader changes in the TME. That these coordinated programs are detectable at the transcriptional level also makes them measurable: using patient-matched imaging and bulk RNA sequencing, we developed a classifier that accurately identifies STaN-high and STaN-low tumors across independent transcriptional cohorts, demonstrating that STaN abundance leaves a reproducible molecular footprint detectable without direct imaging. Given the growing availability of bulk transcriptomic datasets, this approach provides a scalable framework for investigating neural remodeling in cohorts lacking multiplex imaging and illustrates more broadly how quantitative spatial phenotypes can be integrated with transcriptomics to infer otherwise inaccessible TME features.

The transcriptional analyses do not, however, identify the cellular interactions that establish or maintain this phenotype. Spatial organization offers a clue: noradrenergic TH-mid and synaptophysin-high STaNs preferentially localized within tumor-adjacent stromal regions rather than being uniformly distributed, suggesting that neural remodeling occurs within discrete stromal niches. Given the well-established role of stromal compartments in extracellular matrix remodeling, inflammatory signaling, and immune regulation, this spatial pattern raises the possibility that interactions between peripheral nerves, immune, and stromal cells contribute to the adverse phenotypes in STaN-high tumors.

To identify candidate cellular mediators of these spatial relationships, we integrated an independent published single-cell RNA-sequencing atlas of bladder urothelial carcinoma. Because this dataset was not generated from the tumors analyzed here, it cannot establish direct cellular interactions within our cohort. Instead, it provides orthogonal evidence identifying stromal populations with the molecular capacity to participate in reciprocal neural signaling. Inflammatory cancer-associated fibroblasts (iCAFs) broadly expressed secreted neurotrophic factors, neural adhesion molecules, and axon-guidance proteins, while myofibroblastic CAFs (myoCAFs) and mast cells emerged as potential intermediaries of sympathetic signaling. The prognostic associations of noradrenergic STaNs may therefore reflect sympathetic communication with stromal and immune populations rather than direct signaling to tumor cells. The combined noradrenergic-responsive and neurotrophic features of myoCAFs are especially suggestive of a reciprocal relationship in which fibroblasts both respond to sympathetic input and support neural recruitment or maintenance.

Given the preferential localization of prognostic STaNs within fibroblast-rich stroma, these findings suggest that fibroblasts provide a permissive microenvironment for neural recruitment and signaling, consistent with an emerging view of CAFs as central organizers of the TME that communicate reciprocally with peripheral nerves during tissue repair and chronic inflammation. Although these interactions remain poorly characterized in BCa and will require functional validation, the convergence of independent spatial and transcriptomic datasets identifies CAFs as compelling candidate intermediaries. Together, these observations raise the possibility that STaNs act through two complementary modes of communication: paracrine signaling via stromal and immune populations, and short-range or contact-dependent interactions with tumor cells. The latter is reinforced by urothelial expression of postsynaptic machinery alongside the preferential localization of SYP-high STaNs within 0-25 μm of the tumor border, raising the possibility of synapse-like or other contact-dependent STaN-tumor interactions.

An important open question is whether neural remodeling drives aggressive tumor behavior or emerges as a consequence of it. The transcriptional enrichment of neurotrophic factors and axon-guidance pathways within STaN-high tumors suggests that aggressive tumors may actively recruit nerves. Conversely, neurotransmitters and neuropeptides are well-established regulators of immune function, stromal activation, angiogenesis, and extracellular matrix remodeling, providing mechanisms by which neural signaling could reinforce tumor progression and therapeutic resistance. These possibilities are not mutually exclusive and likely reflect dynamic, reciprocal interactions among tumor cells, CAFs, immune populations, and peripheral nerves. Resolving the temporal sequence and functional consequences of these interactions will require mechanistic studies in genetically tractable experimental models with perturbation of candidate neural signaling pathways.

Several limitations warrant consideration. This study was observational and cannot establish causality between neural remodeling and disease progression; functional validation is needed to determine whether peripheral nerves actively drive these processes or respond to broader microenvironmental remodeling. Healthy bladder tissue was not available for comparison, limiting our ability to define how the STaN landscape differs from normal bladder innervation. STaN phenotyping was constrained by the available marker panel, and as a result, the molecular identity of cluster 7 remains unresolved. Technical challenges inherent to imaging peripheral nerve fibers, including discontinuous marker expression, the three-dimensional architecture of neural processes, and the presence of mixed sensory-autonomic nerve bundles, may influence segmentation and phenotype assignment. Finally, the single-cell RNA-sequencing atlas used to nominate candidate stromal interactions was generated from an independent cohort lacking matched multiplex imaging and should be interpreted as complementary evidence rather than direct validation.

Together, this study provides, to our knowledge, the most comprehensive characterization of stromal tumor-associated nerves in human NMIBC to date, establishing STaNs as reproducible microenvironmental features and as a framework for investigating neural remodeling in human cancers, including candidate STaN-stromal, neuroimmune, and STaN-tumor interactions that warrant future mechanistic investigation. More broadly, this work illustrates the value of pairing quantitative spatial phenotyping with transcriptomics to uncover features of the tumor microenvironment that neither modality captures alone.

## METHODS

### Study cohort

The final high-risk, treatment-naïve NMIBC imaging cohort comprised 151 patients (**Table 1**). The median age at diagnosis was 70 years (range=37-88). Female patients represented 11.2% of the cohort with known sex (53% male; 35.8% unannotated). As reported in de Jong et al.^45^ and previously published^64^, tumors were graded by an expert uropathologist according to WHO 2016 criteria^65^. Patients were only included if pathology review confirmed high-grade (HG) urothelial carcinoma without detrusor muscle invasion at primary TURBT or re-TURBT, including predominantly high-grade T1 tumors with a limited number of high-grade Ta cases (n=141 T1HG; n=9 TaHG). 24 of these cases had concomitant carcinoma *in situ* (CIS).

As described in de Jong et al.^45^, comprehensive clinical annotation included treatment and outcome data following intravesical BCG, where BCG response was defined as absence of high-grade (HG) disease for 6 months following adequate BCG exposure, as per international recommendations^19^. Non-response to BCG (BCG failure) was defined as the development of one of the following: i) biopsy-proven muscle-invasive bladder cancer, ii) persistent T1HG NMIBC after BCG induction, iii) high-grade NMIBC after adequate BCG therapy. Adequate BCG therapy was defined as at least 5 out of 6 BCG induction instillations and at least 2 out of 3 BCG maintenance instillation cycles (median number of instillations=21). Longitudinal outcomes included high-grade recurrence-free survival (HG-RFS; defined as time from TURBT to first high-grade recurrence or censoring) and progression-free survival (PFS; defined as time to progression to muscle-invasive disease ≥T2 or metastasis). Clinical outcome and survival comparisons were restricted to patients who received adequate BCG therapy.

Additional clinicopathologic variables included smoking status (yes/no; n=35 no, n=56 yes, n=60 unknown), T1 substage^64^ (T1-microinvasive (T1m; n=21), T1-extensively invasive (T1e; n=73), or unknown n=57)), tumor focality (unifocal/multifocal/missing; n=71 unifocal, n=77 multifocal, n=3 unknown), and size (small <3 cm vs large ≥3 cm or unknown; n=28 small, n=21 large, n=102 unknown), presence lymphovascular invasion (yes/no/unknown; n=5 yes, n=89 no, n=57 unknown), and presence of concomitant CIS (present/absent; n=24 present, n=127 absent). Molecular characterization from de Jong et al.^45^ included EAU^23^ and EORTC^46^ clinicopathological risk classifications and transcriptomic assignment to established NMIBC molecular subtypes, including UROMOL^49^, BCG response subtype (BRS)^45^, Lund^48^, and Chicago T1BC^50^, along with targeted genomic alterations such as FGFR3.

Representative regions from the same formalin-fixed, paraffin-embedded (FFPE) NMIBC tumor blocks used for RNA isolation were cored and assembled into tissue microarrays (TMAs). Four-micrometer TMA sections were used for multispectral imaging.

### Spatial proteomics and STaN quantification

FFPE tumor tissue sections were subjected to multiplex immunofluorescence staining in collaboration with the Human Immune Monitoring Shared Resource at the University of Colorado Cancer Center. Slides were baked at 60°C for 30 minutes and manually deparaffinized through a xylene gradient prior to automated staining on a BOND RX platform (Leica Biosystems). Multiplex staining was performed by sequential rounds of antigen retrieval, blocking, primary antibody incubation, horseradish peroxidase (HRP)-conjugated secondary antibody detection, and deposition of HRP-reactive Opal fluorophores. Antibodies were stripped between successive staining cycles by heat treatment in antigen retrieval buffer.

The eight-marker panel was stained sequentially in the following order: vesicular glutamate transporter-1 (VGLUT1), Substance P (SP), cluster of differentiation-31 (CD31), protein gene product 9.5 (PGP9.5), vesicular acetylcholine transporter (VAChT), synaptophysin (SYP), tyrosine hydroxylase (TH), and pan-cytokeratin. VGLUT1 (Synaptic Systems, 135302; 1:100) was detected with Opal 570 (1:150); Substance P (Abcam, ab133240, clone EPR3959; 1:100) with Opal 690 (1:150); CD31 (Abcam, ab182981, clone EPR17259; 1:100) with Opal 520 (1:100); PGP9.5 (Proteintech, 14730-1-AP; 1:400) with Opal 480 (1:300); VAChT (Synaptic Systems, 139103; 1:100) with Opal 540 (1:150); synaptophysin (Cell Marque, 760-4595, clone MRQ-40; ready-to-use) with Opal 650 (1:150); TH (Sigma-Aldrich, AB152; 1:200) with Opal 620 (1:150); and pan-cytokeratin (Dako, M3515, clone AE1/AE3; 1:250) with Opal 780 (1:25). TH, VAChT, SP, and VGLUT1 antibodies were diluted in with Renaissance Background Reducing Diluent (Biocare Medical, PD905) to reduce non-specific staining.

Antigen retrieval was performed at pH 9 for VGLUT1, Substance P, CD31, PGP9.5, and VAChT staining cycles and at pH 6 for the synaptophysin, TH, and pan-cytokeratin cycles. An extended (40-min) antigen retrieval cycle was performed before PGP9.5 staining to eliminate residual CD31 signal observed after the preceding staining cycle; standard antigen retrieval conditions were sufficient for all other markers. Opal fluorophores were diluted in 1x Plus Automation Amplification Diluent (Quanterix, FP1609), and detection used 1x Opal Anti-Mouse + Rabbit HRP (Quanterix, ARH1001EA). Spectral DAPI (Quanterix, FP1490) was used for nuclear counterstaining.

Whole-slide multispectral images were acquired using the PhenoImager HT instrument (formerly Vectra Polaris, Quanterix) with a 20x objective at 0.5-µm resolution using PhenoImager HT software v2.1.0. Regions of interest were selected using PhenoChart software v2.2.0 and rescanned using the 20x objective and the multispectral imaging cube. Spectral references and unstained control images were measured and inForm software v3.0 was used to create a multispectral library reference. The multispectral images (.im3 files) were spectrally unmixed and analyzed with tissue-, cell-, and object segmentation using inForm software v3.0 (Quanterix).

Tumor, stromal, glass, lumen, and erythrocytic compartments were defined using a semi-supervised adaptive tissue segmentation algorithm trained on representative user-defined regions using InForm. Within the stromal compartment, threshold-based object segmentation was used to identify and quantify discrete PGP9.5+ and/or SYP+ nerve (STaN) and CD31+ vascular “objects”. Images were visually inspected to identify cores with poor tissue integrity, spectral unmixing, or tissue/nerve segmentation; these images were excluded from the final cohort.

### STaN identification and phenotyping

PGP9.5+/SYP+/-objects from all images were pooled and subjected to K-means clustering based on mean pixel intensity to remove imaging artifacts and other non-neuronal signals (k=3), including autofluorescent erythrocytes affecting the Opal 480 channel and an unexpected non-neuronal population of SYP+/CD31+ cells. Phenotypes were assigned to filtered STaN objects by K-means clustering based on mean pixel intensity. The optimal number of nerve clusters (k=7) was selected based on silhouette scores and within-cluster sum of squares, which supported six to seven clusters. Seven clusters were retained because they provided additional resolution of the SP+/VGLUT1+ nerve population. Image-level nerve features were aggregated to the patient level by summing stromal and STaN+ area across all cores. To account for variability in tissue composition, stromal nerve abundance was defined as the percentage of total STaN+ area relative to total stromal area across all TMA cores per patient. TMA cores with <5% tumor area, insufficient stromal area analyzed (<250,000 µm^2^, approximately one-third of a TMA core), or lacking clinical data were excluded from downstream analyses. All image processing, quality control, segmentation, and filtering were performed blinded to patient-level clinical information and outcomes, which were linked to imaging data only after generation of patient-level summaries.

### Distance-based spatial enrichment analysis

Adaptive tissue segmentation and threshold-based object segmentation were performed using the Akoya Biosciences InForm analysis platform (version 3.0.0) to generate tissue-class segmentation masks for tumor, stroma, and glass and instance-labeled masks for nerves (PGP9.5+/SYP+/−) and vessels (CD31+), as described above. These outputs were converted to single-channel masks for downstream analysis. STaN cluster assignments were mapped to nerve-instance labels using matching image and object identifiers. Downstream image processing for spatial analyses was performed on the University of Colorado Boulder Alpine high-performance computing cluster^66^ using R version 4.4.1 within an RStudio container running Ubuntu 22.04.5 LTS. Image processing used EBImage^67^ version 4.48.0 and fftwtools^68^ version 0.9-11, dynamically linked to FFTW^69^ version 3.3.8 (Ubuntu package libfftw3-double3, version 3.3.8-2ubuntu8). Because convolution-based tissue-component filtering was sensitive to the numerical environment, the software versions, analysis configuration, and computational environment records are provided with the analysis code.

Tumor, stromal and vessel masks were refined using morphological filtering to remove segmentation artifacts and tissue-edge debris while preserving contiguous tissue regions and vascular-like structures. Morphological characteristics were calculated using EBImage and mmand^70^ (R package version 1.6.3). To evaluate spatial organization among tumors with appreciable innervation, spatial analyses were restricted to images from STaN-high tumors (upper tertile of final imaging cohort). Tissue cores with large tears, or less than 250,000 µm² of stroma were excluded. The final spatial analysis cohort included images from 45 TMA cores from 32 patients (BCG response: n=18 BCG failure and n=14 BCG responders; progression: n=12 progression and n=20 no progression).

Images were analyzed at a spatial calibration of 0.5 µm per pixel. The tumor-stroma border was defined as cleaned tumor pixels overlapping a dilation of the cleaned stromal mask, with glass pixels excluded. Dilation was performed using ‘EBImage::dilate()’ with a disc structuring element generated by ‘EBImage::makeBrush(size = 3, shape = “disc”)’. Euclidean distance from the tumor-stroma border was calculated using ‘EBImage::distmap()’ after subsampling the interface mask twofold along each image axis. Distances were converted to micrometers based on the resulting 1 µm grid spacing and resampled to the original image dimensions by nearest-index resampling. Within the cleaned stromal compartment, stromal area and positive area for each of the six STaN clusters and for vessels were quantified across five concentric distance bands from the tumor-stroma border: 0-25, 26-50, 51-100, 101-200, and 201-300 µm. Positive area was calculated from pixels overlapping the cleaned stromal mask, and band-level density was defined as STaN subtype- or vessel-positive area divided by stromal area within the corresponding band. Pixels >300 µm from the tumor-stroma border were excluded from these measurements. Images without sufficient stromal tissue to extend to the 300 µm distance band were excluded from downstream spatial modeling.

Nerve and vascular spatial enrichment within concentric stromal distance bands from the tumor-stroma interface was evaluated at the image level using Tweedie generalized linear mixed models implemented with the glmmTMB^71^ R package (version 1.1.14). Separate models were fit for each nerve subtype and for vasculature, with the percentage of stromal band area occupied by nerve or vascular signal as the response and a Tweedie distribution with log link to accommodate zero-containing, non-negative, right-skewed measurements. Because STaN subtype composition varied across images and patients, subtype-specific models included only images containing the corresponding STaN cluster. STaN clusters were omitted if fewer than n=10 patients per outcome contained that STaN subtype; this led to exclusion of STaN cluster 1 (SP+/VGLUT1+) from modeling. Models included fixed effects for distance band, clinical outcome, and their interaction, with the natural logarithm of the overall nerve or vascular area fraction within the full analyzed stromal radius included as an offset, thereby evaluating band-specific occupancy relative to overall regional abundance. Patient and image identity were modeled as nested random intercepts to account for multiple distance-band measurements within images and multiple images per patient. To account for differences in tissue area within stromal bands and nerve or vascular representation across images, observations were weighted by the product of the square roots of stromal area within each band and total nerve or vascular area within the analyzed radius; nerve or vascular area used for weighting was capped at the 99th percentile, and weights were normalized to a mean of one. Estimated marginal means and contrasts were computed using the emmeans^72^ R package (version 1.11.1). Within each outcome group, enrichment in each distance band was compared with the equally weighted mean of all remaining bands; outcome groups were additionally compared within each distance band, and differences in band-versus-rest enrichment between outcomes were tested. P values were adjusted for multiple comparisons within each STaN subtype or vascular analysis using the Benjamini-Hochberg method. Enrichment differences were visualized using split bubble plots generated with custom scripts adapted from visualization approaches implemented by Canete et al. in the spicyR^73^ R package.

### Object-to-border spatial distance analysis

Vascular objects and objects of each STaN subtype were identified by connected-component labeling of their respective binary masks. Object boundary contours were extracted as ordered coordinate sequences using EBImage::ocontour(), representing each object by its boundary rather than by a single centroid. The tumor–stroma border was defined as described above and decomposed into connected segments. For each segment, a Euclidean distance transform was calculated using EBImage::distmap() after subsampling the border-segment mask by a factor of two along each image axis. Distances were converted to micrometers using the resulting 1 µm grid spacing, corresponding to an original resolution of 0.5 µm per pixel, and resampled to the original image dimensions using nearest-index resampling. Each STaN or vascular object’s distance to the tumor-stroma border was defined as the minimum distance across its boundary coordinates and all border segments, providing an edge-to-border rather than centroid-to-border measurement. Distributions of STaN and vascular distances to the tumor-stroma border were visualized as ridgeline plots using the ggridges R package (version 0.5.6).

### Survival analyses

Kaplan-Meier survival estimates for high-grade recurrence-free and progression-free survival were performed using the survival^74^ R package (version 3.8-3). Patients were stratified by median or upper/lower tertiles of STaN abundance, NMIBC subtype, combined NMIBC subtype and STaN abundance, or predicted STaN status as indicated in the figure legends. Survival differences were assessed using the log-rank test. Kaplan-Meier plots were visualized with the survminer^75^ R package (version 0.5.0).

Multivariable Cox proportional hazards models were used to evaluate associations between stromal nerve abundance, clinicopathological features, molecular subtype, and clinical outcomes. Hazard ratios (HRs) and 95% confidence intervals (CIs) were reported. Multivariable Cox proportional hazards models were constructed adjusting for relevant clinical covariates with n>5 per STaN-status and outcome group and <10% missing/unknown. Given the homogeneity of the cohort regarding stage and grade, and substantial missingness within the clinical annotations, the final model was only adjusted for age. Cox model results were visualized using forest plots generated using custom scripts with the ggplot2 R package (version 4.0.3).

### Bulk gene expression normalization and filtering

Raw counts from BRS Cohorts A and B were obtained from the BRSpred^45^ R/Bioconductor package (version 1.2.1). Cohorts A and B were sequenced on different platforms, as previously described^45^. Raw counts were filtered to remove lowly expressed genes, retaining those with ≥8 counts in at least five patients, and were subsequently converted to log_2_-transformed counts per million (CPM) without TMM normalization using edgeR^76^ (version 4.0.16). Raw counts were transformed to log_2_ counts per million (log_2_-CPM) with a prior pseudocount of 1 for downstream gene set enrichment analyses and machine learning.

Gene annotations were obtained from BioMart (Ensembl Genes 116; GRCh38.p14). Unless otherwise specified, genes were filtered by biotype to include protein-coding genes, long non-coding RNAs (lncRNAs), and microRNAs (miRNAs). The resulting log_2_-CPM expression matrices were used as input for elastic net model training and validation. All normalization and filtering steps were performed independently across cohorts and across data subsets within each cohort (training and unseen subsets of cohort A; test and unseen subsets of cohort B).

### Differential gene expression and functional gene set enrichment

Differential expression analysis between STaN-high (upper tertile; n=33) and STaN-low (lower tertile; n=33) tumors was performed on raw counts using DESeq2^51^ (version 1.42.1). Cohort and sex were included as covariates to correct cohort-specific batch effects and sexual dimorphism, respectively. Genes with an adjusted p-value (Benjamini-Hochberg) < 0.05 and absolute log_2_-fold change (|log_2_-FC|) > 1 were considered significantly differentially expressed. Results were visualized using volcano plots and heatmaps of the top differentially expressed genes (ranked by adjusted p-value and/or log_2_-FC). Gene set enrichment analysis (GSEA) was performed using the clusterProfiler^77^ R package (version 4.10.1) with gene sets from the Hallmark (H), Kegg 2026 Canonical Pathways, and Gene Ontology Biological Processes (C5 GOBP) MSigDB collection(s) and PanglaoDB cell type signature database. Gene sets were obtained from the enrichR R package (version 3.4). Genes were ranked by Wald statistic for pre-ranked GSEA. To reduce redundancy among enriched pathways for visualization, pairwise overlap between gene sets was quantified using the Jaccard index of leading-edge genes, and highly overlapping terms were collapsed using a similarity threshold of 0.25, retaining representative pathways based on normalized enrichment score (NES) and adjusted p-value. GSEA results were visualized by dot plots generated using custom R scripts. Gene sets were considered significantly enriched when |NES| > 1 and FDR < 0.05. DEseq2 and raw GSEA results are provided in **Table S2**.

### Training and evaluating the STaN-high/low classifier

log_2_-CPM gene expression data from BRS Cohort A were used for model training, while BRS Cohort B served as an independent test cohort that was not used during model development (**Table S3**). As described in de Jong et al.^45^, Cohort B was sequenced on a different platform from Cohort A, providing an additional source of technical independence for model evaluation. Among patients with matched STaN quantification and gene expression data, STaN-high and STaN-low tumors were defined as the upper and lower tertiles of stromal nerve area. To increase training and test cohort sizes and balance STaN classes within each cohort, STaN tertiles were calculated separately for each cohort. Patients in BRS Cohort A (train; n=23 STaN-high, n=23 STaN-low) were used for feature selection, hyperparameter tuning, and final model training. Patients in BRS Cohort B (test; n=10 STaN-high, n=10 STaN-low) were used as an independent validation cohort to evaluate model performance.

Feature selection and elastic net modeling were performed using glmnet^78^ (version 4.1-9) to predict STaN class (STaN-high versus STaN-low tumors) as a binary variable. The top 1,000 most variable genes, ranked by variance across the training dataset, were retained for model development. Feature stability was assessed across 1,000 bootstrap resamples of the training dataset. Models were fit to each bootstrap resample using a fixed elastic net mixing parameter (alpha = 0.5) and the 1-standard-error regularization parameter (lambda.1se) obtained from 10-fold cross-validation of the full training dataset. Genes were retained for downstream modeling if they had non-zero coefficients in at least 25% of bootstrap models and maintained the same coefficient direction in at least 75% of models in which they were selected.

Hyperparameter tuning was performed using 10-fold cross-validation with shared folds across candidate alpha values ranging from 0 to 1.0 in increments of 0.1. Because the feature space was stringently reduced by bootstrap stability selection using lambda.1se, lambda.min was used during final model tuning to minimize cross-validated deviance without imposing additional feature sparsity. For each alpha, the lambda value minimizing cross-validated deviance (lambda.min) was identified. The overall minimum cross-validated deviance and its standard error were used to define a 1-standard-error threshold across alpha values, and the largest alpha whose minimum deviance fell within this threshold was selected. The final classifier was fit to the full training dataset using the selected alpha and its corresponding lambda.min (alpha = 0.3; lambda = 0.0552196).

Independent validation was performed using BRS Cohort B (test), which was not used for feature selection, hyperparameter tuning, or model fitting. Model performance was evaluated using area under the receiver operating characteristic curve (AUC), sensitivity, specificity, and accuracy. Receiver operating characteristic (ROC) curves were generated using the pROC^79^ R package (version 1.18.5), and confusion matrices were generated using caret (version 7.0-1).

### Clinical validation of classifier predictions

The STaN-high/low classifier was used to predict stromal nerve abundance in unseen patients from BRS Cohorts A and B that were not included in multispectral imaging (n=50 cohort A, n=110 cohort B) and therefore lack image-based nerve quantification. Similarly, we applied the classifier to the patients who received BCG in two independent external NMIBC cohorts UROMOL^49^ and Robertson et al. T1 high-grade NMIBC (Chicago T1BC)^50^). Model predictions (STaN-high vs STaN-low) were used to stratify patients for Kaplan-Meier estimates of high-grade recurrence-free and progression-free survival.

### Single-cell RNA-seq integration, clustering, and annotation

A previously published single-cell RNA-sequencing atlas of human bladder urothelial carcinoma from Chen et al.^52^ was obtained from NCBI Sequence Read Archive (SRA) under project accession number (PRJNA662018) using the SRA Toolkit. Raw single-cell RNA-sequencing data from three normal and eight bladder tumor samples were aligned with Cell Ranger, and the resulting filtered feature-barcode matrices were imported into R (v4.5.0) using Seurat^80^ (v5.0.0) via ‘Seurat::Read10X()’. Cells with 300-6,000 detected genes, >1,000 UMIs, and <10% mitochondrial reads were retained, and doublets were removed per sample using DoubletFinder (v2.0). Data were merged and normalized with ‘Seurat::SCTransform()’, regressing out mitochondrial percentage and cell cycle scores. PCA was performed, retaining PCs up to the point of diminishing variance explained (elbow/cumulative variance method), and batch effects were corrected by reciprocal PCA (RPCA) integration. Clustering was performed on the RPCA-integrated embedding using a shared nearest-neighbor graph and the Louvain algorithm (resolution = 0.1), and visualized by UMAP. Clusters were manually annotated into eleven cell types (Urothelial cell, T cell, Endothelial, MyoCAF, iCAF, Myeloid, Proliferating cell, B cell, Differentiating plasma cell, Mast cell, and Mature plasma cell) based on expression of canonical marker genes.

For downstream analyses, analyses relevant to NMIBC were restricted to cells from high-grade non-muscle-invasive tumors (n=2). Genes identified as differentially expressed between STaN-high and STaN-low tumors in the bulk RNA-sequencing analysis on the BRS cohort were mapped onto the single-cell dataset to determine their distribution across annotated cell populations. Genes meeting |log_2_-FC| ≥ 1.5 and adjusted P ≤ 0.05 in the bulk analysis and detected in the single-cell dataset were retained. Mean expression across cell populations was calculated using ‘Seurat::AverageExpression()’ from the SCT assay, followed by gene-wise z-score transformation for visualization.

For heatmap visualization, genes and cell populations were hierarchically clustered using Pearson correlation distance and Ward’s D2 linkage, and genes were partitioned into expression modules using repeated k-means clustering. In complementary analyses of candidate neural signaling programs, curated genes encoding neurotrophic factors, neurotransmitter receptors, neural adhesion molecules, axon-guidance molecules, and related signaling components were examined across annotated cell populations. Mean normalized RNA expression for these analyses was calculated from the RNA assay using ‘Seurat::AverageExpression()’. Heatmaps of differentially expressed and proposed neuro-stromal communication network genes were visualized using the ComplexHeatmap R package (version 2.18.0).

### Statistical analyses

Unless otherwise stated, all pairwise comparisons were performed using the Wilcoxon rank sum non-parametric test. P-values were adjusted for multiple hypothesis testing using the Benjamini-Hochberg method. Adjusted p-values were reported as false discovery rate (FDR). Baseline characteristics were compared across STaN tertiles using Fisher’s exact test for categorical variables and the Kruskal-Wallis test for age. Patients with missing data for a given variable were excluded from the corresponding analysis. Unless otherwise stated, all statistical tests were performed in R (version 4.3.3).

## Data and code availability

Raw multispectral images (.im3 files), spectrally unmixed component images, binary tissue, nerve, and vessel masks, processed spatial masks used for distance-based analyses, and derived nerve-, and image-level quantitative data are available under the BioImage Archive (http://www.ebi.ac.uk/bioimage-archive) under accession S-BIAD-4129. Patient-level quantitative data and clinical annotations are available in **Table S1**. Raw sequencing data for BRS Cohorts A and B is available for public use in the European Genome-Phenome Archive under accession number: EGAS00001006879. Count-level matrices are available through the BRSpred^45^ R package (version 1.2.1). Code used for image processing, spatial, clinical, survival, bulk transcriptomic, and single-cell transcriptomic analyses is available at https://github.com/cdeit/NMIBC_innervation. Differentially expressed genes and raw GSEA results are available in **Table S2**. All code and data to reproduce, evaluate, or apply the TME STaN-high/low classifier are available through the gotNeRve R package at https://github.com/cdeit/gotNeRve. Training and evaluation expression matrices, model coefficients, selected features, and model predictions are available in **Table S3**. Phenotyped nerve-level data matrices are available in **Table S4**. Spatial metrics and modeling statistics are available in **Table S5**. The single cell transcriptomic dataset from Chen et al.^52^ is available in GSA-Human under the accession code HRA000212 and in SRA datasets under BioProject PRJNA662018. Raw bulk transcriptomic data from the Robertson et al. Chicago T1BC^50^ external validation cohort is available through the Gene Expression Omnibus (GEO) under accession code GSE154261. Raw bulk transcriptomic data for the UROMOL^49^ external validation cohort are available under controlled access through the European Genome-phenome Archive (EGA) under accession code EGAS00001004693.

## Acknowledgements

CSD thanks Ralf P Dagdag, Lily Elizabeth R Feldman, Lucas E Gillenwater, Michael V Orman, Sutanu Nandi, Kyle Sullivan, and Natalie Zajczenko for their support and insightful discussions throughout this work. We thank the patients and their families whose participation made this research possible. We thank Troy Schedin, BS, HT, and the Human Immune Monitoring Shared Resource (RRID:SCR_021985) at the University of Colorado Anschutz Medical Campus for technical assistance with multiplex immunofluorescence staining and imaging.

## Funding

This work was generously supported by the Cancer League of Colorado (AWD-242959) to JCC, the Predoctoral Training Grant in CU Anschutz Pharmacology and Molecular Medicine (T32GM158468) to CSD, an NIH/NCI supplement to R01CA268055 to JCC supporting CSD, MRACE (107477) to TZ, and NIH/NCI R50CA293845 to KRJ. This work was supported in part by the University of Colorado Cancer Center Support Grant (P30CA046934). This work utilized the Alpine High-Performance Computing resource at the University of Colorado Boulder. Alpine is jointly funded by the University of Colorado Boulder, the University of Colorado Anschutz, Colorado State University, and the National Science Foundation (award 2201538).

## Competing Interests

JCC is a cofounder and Chief Science Officer of OncoRx Insights. The other authors declare no competing interests.

## Declaration of generative AI and AI-assisted technologies in the manuscript preparation process

During the preparation of this work, the authors used ChatGPT (OpenAI) and Claude (Anthropic) for editing, reviewing, and checking the manuscript for inconsistencies and errors. The authors reviewed and edited the output as needed and take full responsibility for the content of the published article.

## STAR methods key resource table

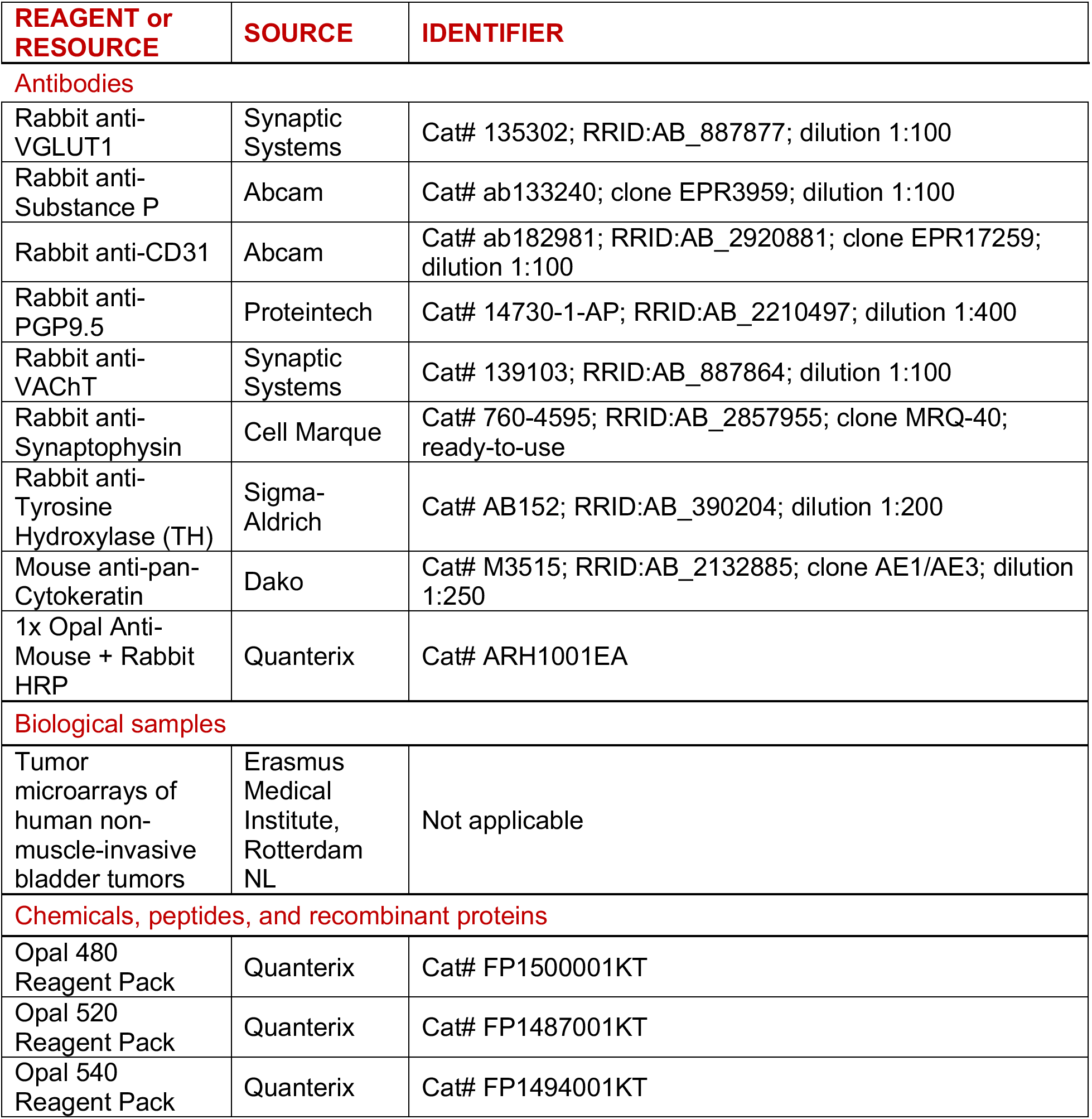
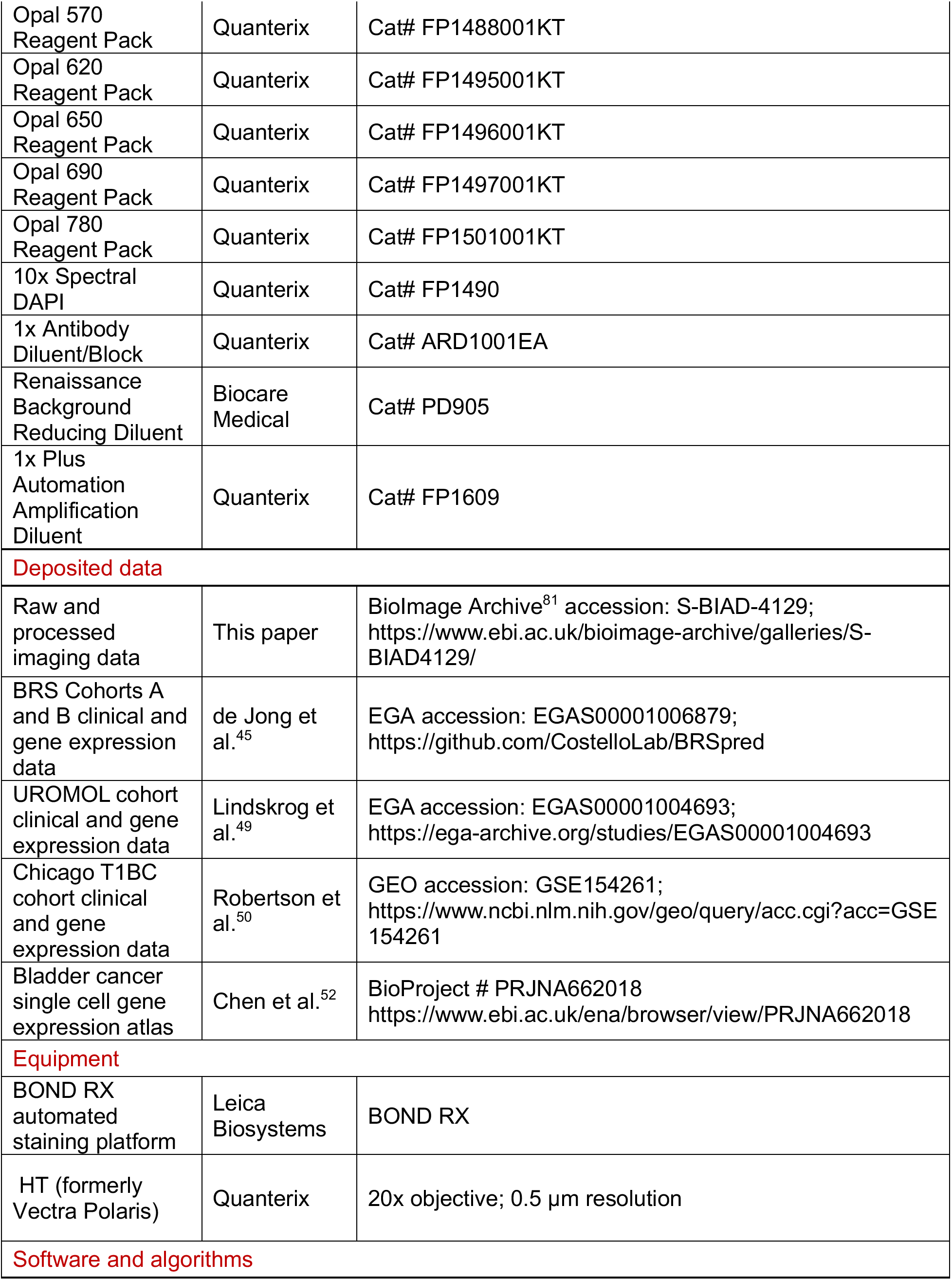
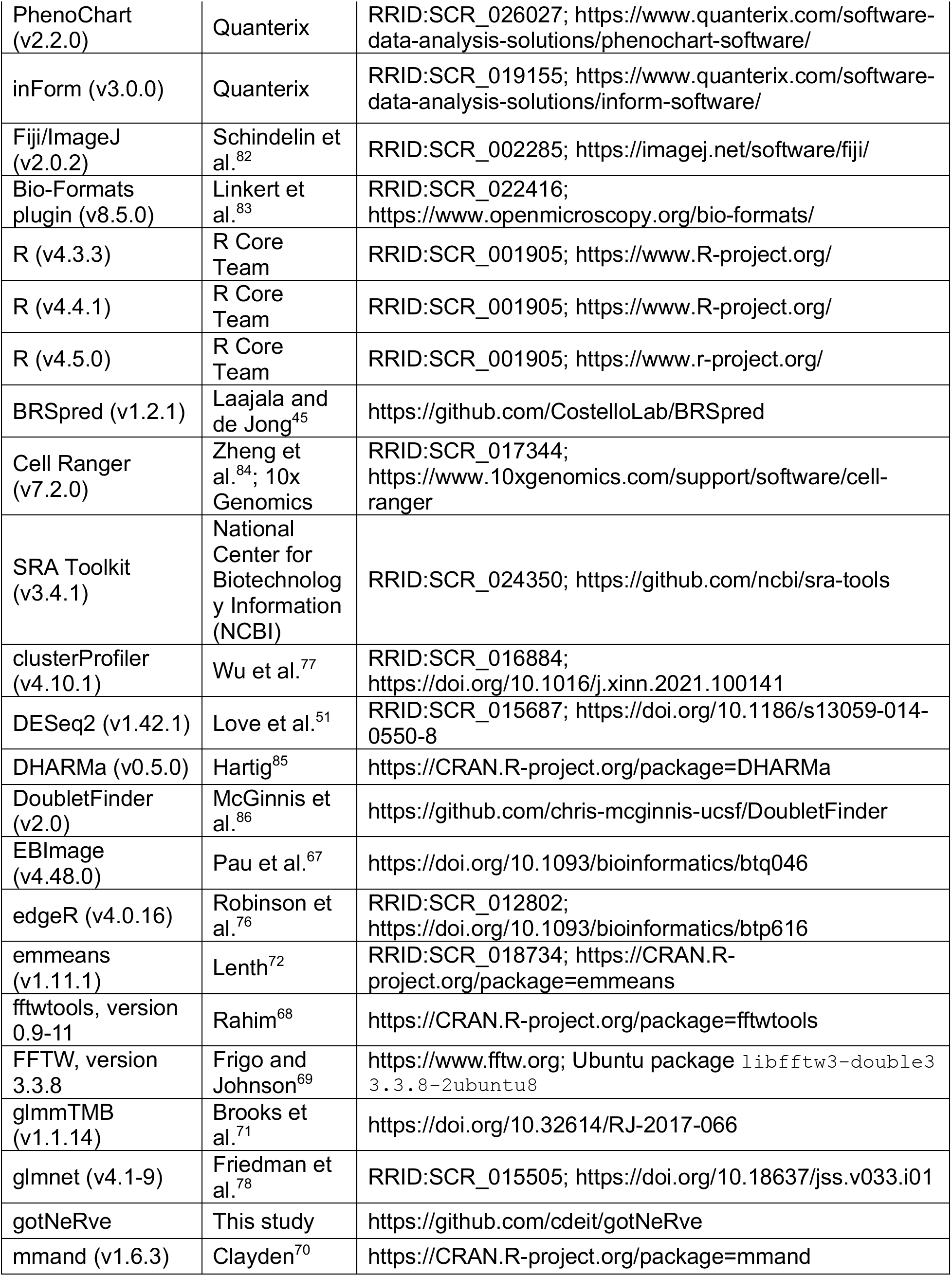
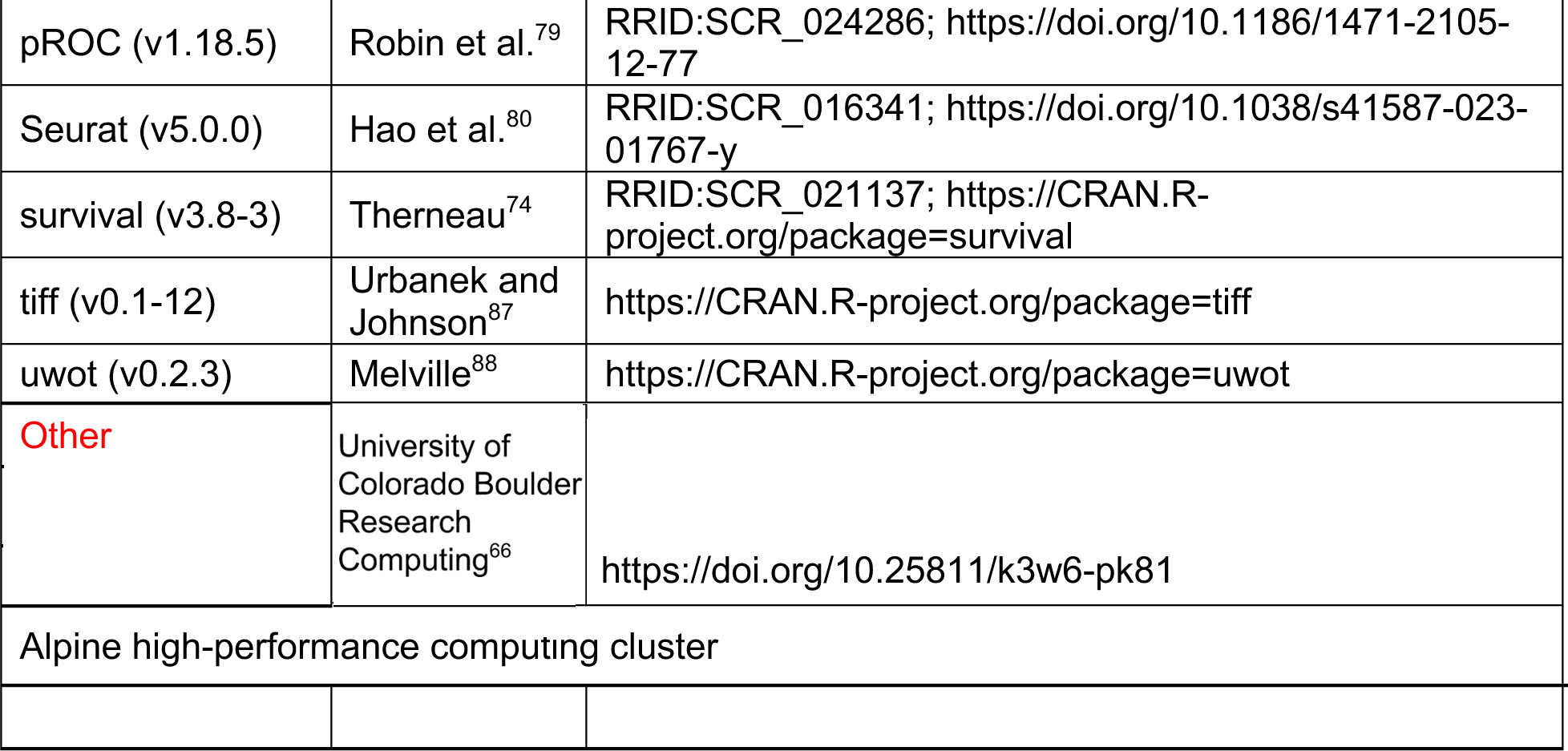

## Notes

https://github.com/cdeit/gotNeRve

https://github.com/cdeit/NMIBC_innervation

### Competing Interest Statement

J.C.C. is a cofounder and Chief Science Officer of OncoRx Insights. The other authors declare no competing interests.

