## Supplementary Figures for "Stromal innervation is prognostic in non-muscle-invasive bladder cancer and predicted by a distinct microenvironmental transcriptomic signature"

**Figure S1.** Associations between STaN abundance and additional clinicopathological characteristics.

**Figure S2.** Survival outcomes stratified by median STaN abundance.

**Figure S3.** STaN abundance by EORTC risk group and Lund BCa subtype class.

**Figure S4.** Functional gene set enrichment analysis of GO biological processes and PanglaoDB cell types in STaN-high vs STaN-low tumors.

**Figure S5.** UMAP embedding of single cell clusters from external scRNA-seq dataset.

**Figure S6.** Development and validation of an elastic net classifier for prediction of STaN status from bulk RNA sequencing.

**Figure S7.** Expression of 12-gene STaN signature genes in STaN-high vs STaN-low tumors.

**Figure S8.** Performance of the transcriptional STaN abundance class classifier and external clinical validation.

**Figure S9.** Predicted STaN class provides additional prognostic stratification within established NMIBC clinicopathological risk groups and molecular subtypes.

**Figure S10.** STaN phenotype classification by k-means clustering.

**Supplementary Tables**

**Table S1. Clinical data and patient-level STaN abundance for the final imaging cohort.**

**Table S2.** Differential gene expression and gene set enrichment analyses associated with STaN abundance. **(A)** Differentially expressed genes between STaN-high and STaN-low tumors. Positive log2 fold-change (log2FC) values indicate higher expression in STaN-high tumors, whereas negative values indicate higher expression in STaN-low tumors. **(B–E)** Raw gene set enrichment analysis (GSEA) results using **(B)** Hallmark, **(C)** KEGG, **(D)** Gene Ontology Biological Process (GOBP), and **(E)** PanglaoDB gene sets. Positive normalized enrichment scores (NES) indicate enrichment in STaN-high tumors, whereas negative NES indicate enrichment in STaN-low tumors.

**Table S3. Cohort overview and feature selection for training and evaluating the elastic net STaN-high/low classifier. (A)** Clinical data and model predictions for the training, test, and unseen clinical validation cohorts. **(B)** Top 1,000 genes by variance in the training cohort. **(C)** Genes retained by bootstrap stability selection. **(D)** Genes with non-zero coefficients in the final model.

**Table S4. Clustered PGP9.5+/SYP+/− objects, candidate STaN objects, and tissue regions.**

**Table S5. Spatial metrics and statistical analyses used for spatial analyses. (A, B)** Tweedie model statistics for spatial enrichment in relation to **(A)** BCG failure and **(B)**

progression. **(C, D)** Spatial metrics for STaN and vessel masks relative to the tumor-stroma border. **(C**) Distances of STaNs and vessels from the tumor-stroma border. **(D)** Spatial enrichment of STaNs and vessels within concentric distance bands from the tumor-stroma border.

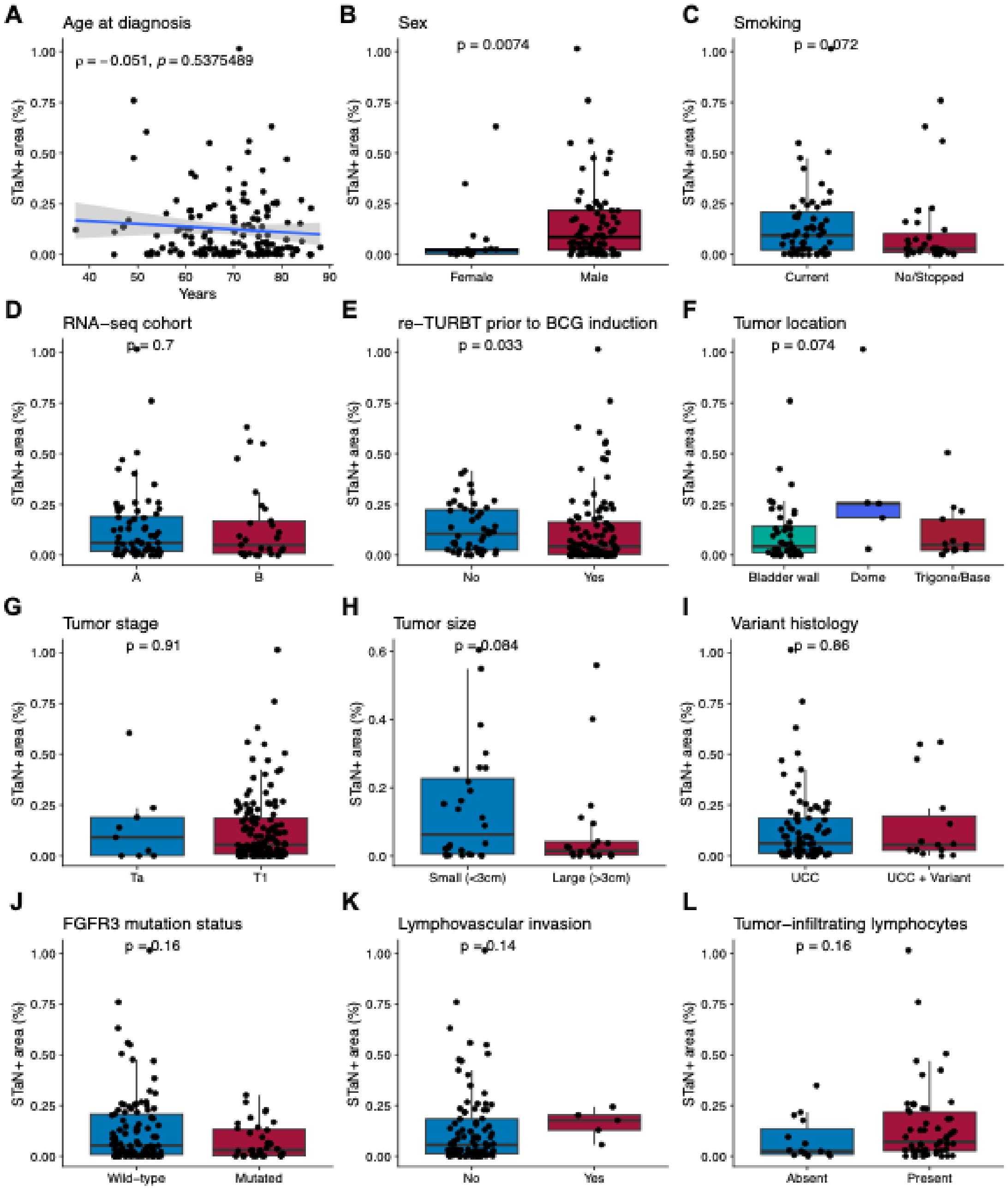


**Supplementary Figure 1. Associations between STaN abundance and additional clinicopathological characteristics.** STaN abundance was evaluated according to **(A)** age at diagnosis, **(B)** sex, **(C)** smoking history, **(D)** RNA-sequencing cohort assignment from the BRS study, **(E)** repeat transurethral resection (re-TURBT) prior to BCG induction, **(F)** tumor location, **(G)** tumor stage, **(H)** tumor size, **(I)** variant histology, **(J)** FGFR3 mutation status, **(K)** lymphovascular invasion, and **(L)** tumor-infiltrating lymphocytes. Points represent individual tumors. The blue line in **(A)** indicates the linear regression fit with the 95% confidence interval. Statistical significance was assessed using Spearman correlation for age, two-sided Wilcoxon rank-sum tests for two-group comparisons, and the Kruskal-Wallis test for tumor location. P values are shown on each panel.

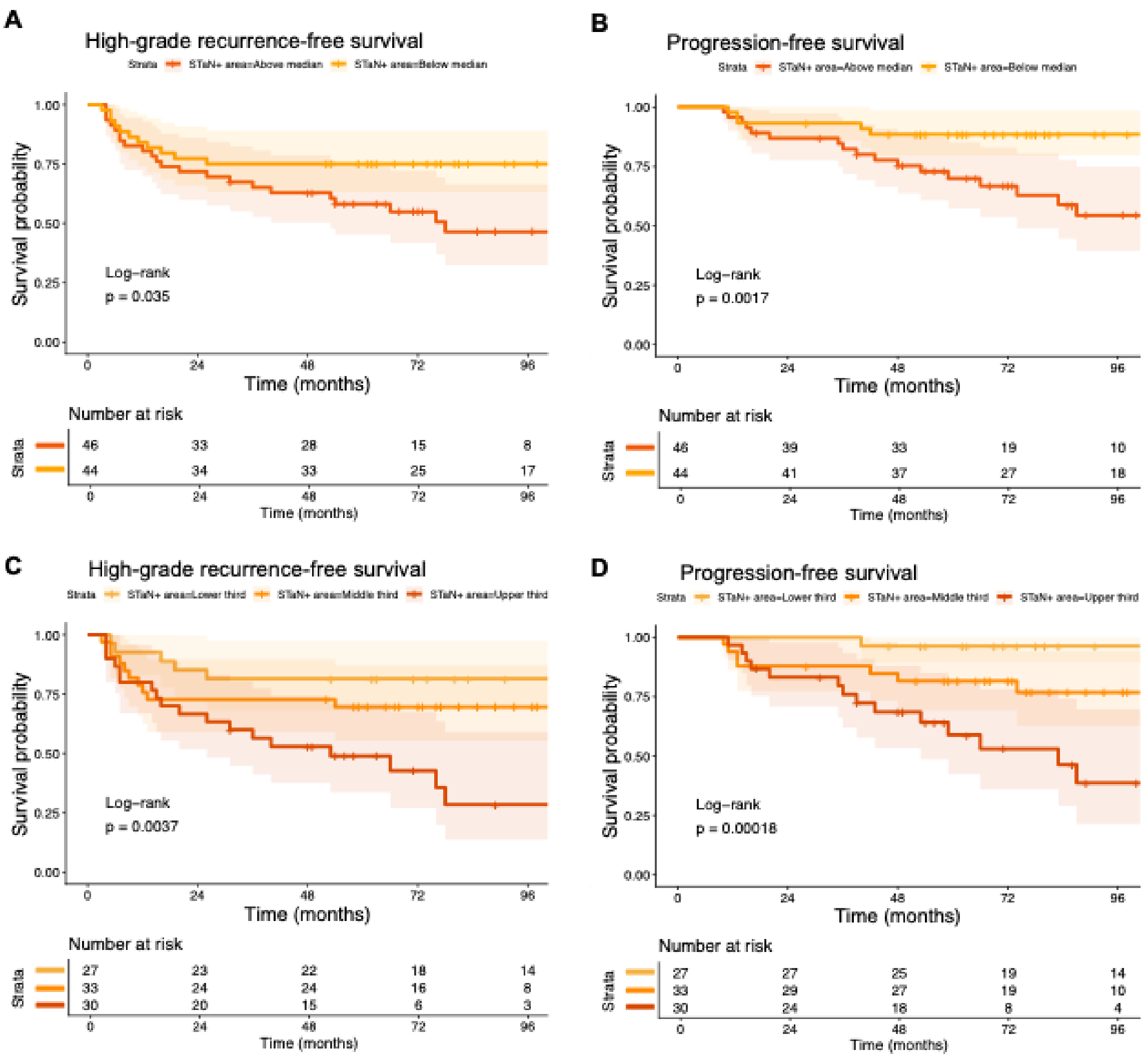


**Supplementary Figure 2. Survival associations with STaN abundance are robust to alternative stratification approaches.** Kaplan-Meier analysis of high-grade recurrence-free survival (HG-RFS) and progression-free survival (PFS) using **(A, B)** median-based stratification and **(C, D)** all three STaN abundance tertiles. Above-median STaN abundance was associated with worse HG-RFS (log-rank, p = 0.035) and PFS (log-rank, p = 0.0017). Survival also differed significantly across STaN tertiles for both HG-RFS (log-rank, p = 0.0037) and PFS (log-rank, p = 0.0018), with the upper tertile exhibiting the poorest outcomes. Shaded regions indicate 95% confidence intervals; tick marks indicate censored observations.

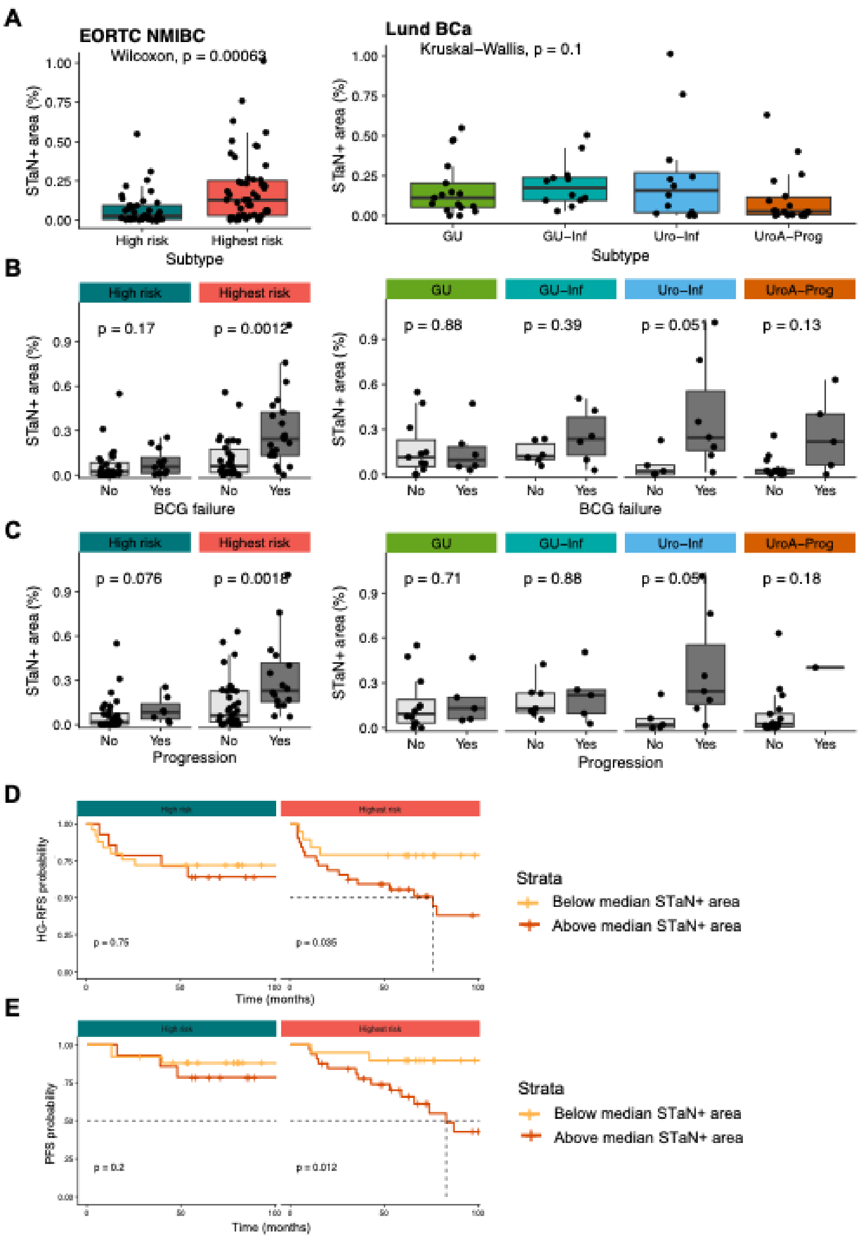


**Supplementary Figure 3. STaN abundance by European Organisation for Research and Treatment of Cancer (EORTC) NMIBC histopathological risk group and Lund BCa molecular subtype. (A)** STaN+ area according to subtype classifications with >=10 patients total and >=3 patients per clinical outcome. Each data point represents a single patient. **(B,C)** STaN+ area according to **(B)** BCG failure status and **(C)** progression status. Boxes indicate the interquartile range (IQR), center lines denote the median, and whiskers extend to1.5x IQR. Each data point represents a single patient. Statistical significance was assessed using the Wilcoxon rank-sum test or Kruskal-Wallis test for groups with more than two comparisons. Adjusted p-values are indicated. Patient STaN abundance is defined as STaN+ area (%) (STaN+ area ($um^2$) / total stroma area ($um^2$) * 100). **(D-E)** Kaplan-Meier survival analyses of high-grade recurrence-free survival (HG-RFS) **(D)** and progression-free survival (PFS) **(E)** stratified by median STaN+ area within each subtype. Median stratification was used in place of upper/lower STaN tertiles to increase sample sizes per group. Statistical significance was assessed using the log-rank test. P-values are indicated.

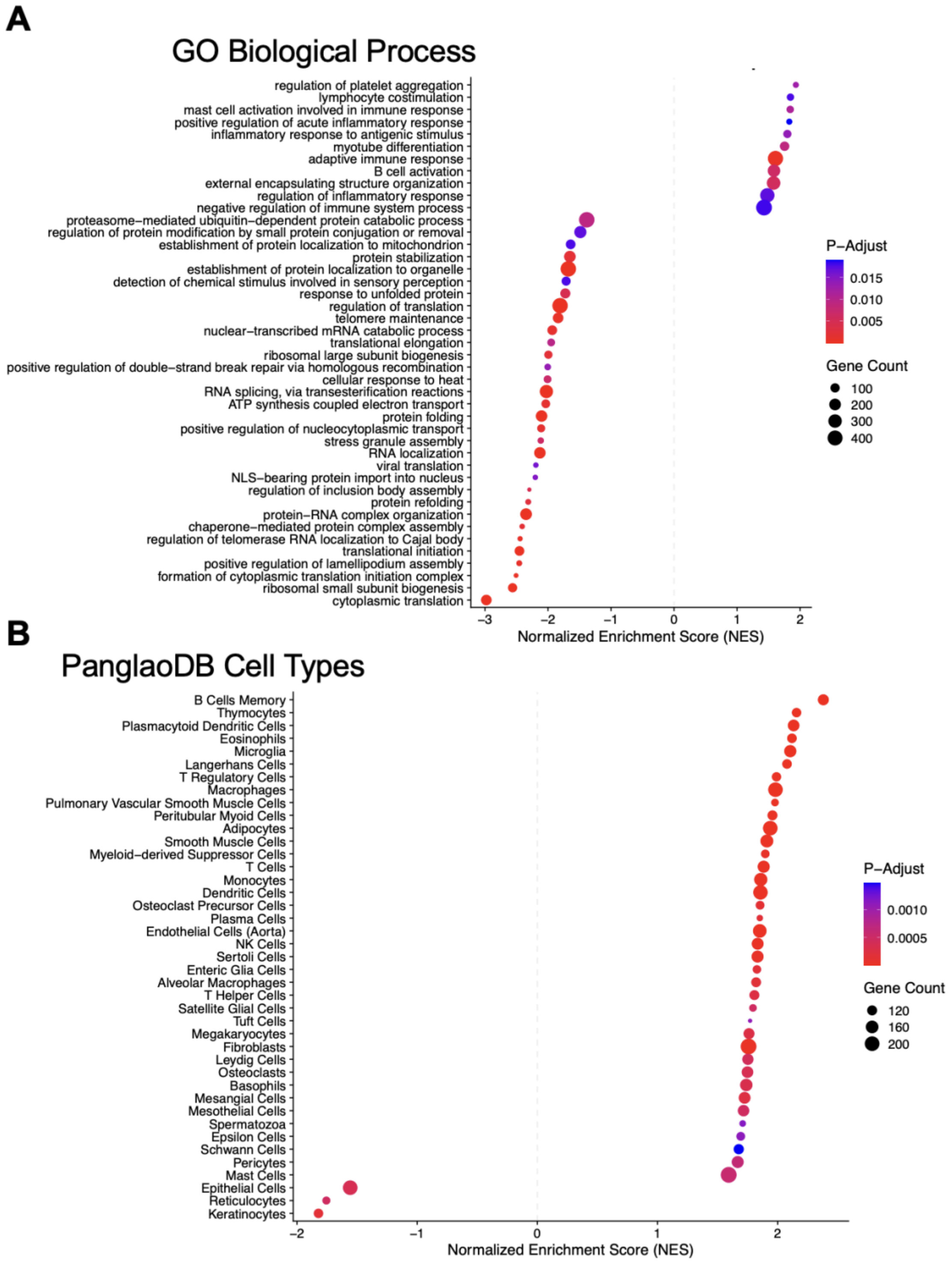


**Supplementary Figure 4. Functional gene set enrichment analysis of GO biological processes and PanglaoDB cell types in STaN-high vs STaN-low tumors. (A)** Gene Set Enrichment Analysis (GSEA) of Gene Ontology (GO) Biological Process terms associated with STaN-high (upper tertile; NES > 1) versus STaN-low (lower tertile; NES < 1) tumors. Redundant GO terms were reduced using semantic similarity-based clustering implemented in clusterProfiler. **(B)** GSEA of PanglaoDB cell type signatures. Redundant cell type signatures were collapsed using Jaccard similarity-based clustering. Bubble size represents the number of genes contributing to each enriched gene set, color indicates the Benjamini-Hochberg adjusted p value, and the x-axis shows the normalized enrichment score (NES).

A

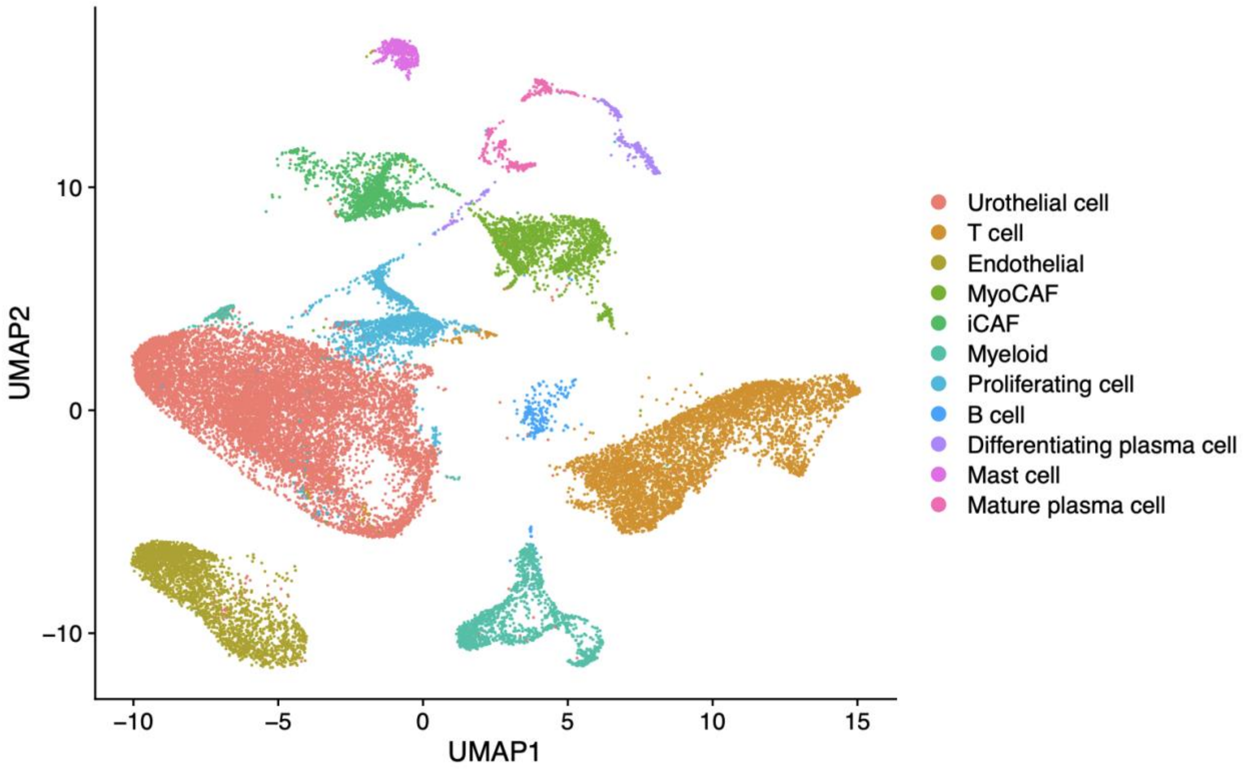


**Supplementary Figure 5.** UMAP embedding of integrated single cell clusters from external scRNA-seq reference (Chen et al. [52]). The dataset was reintegrated, reclustered, and annotated to define major cell populations. The resulting reference atlas was used to identify the putative cellular sources of STaN-associated differentially expressed genes and neuro-stroma-immune interaction network.

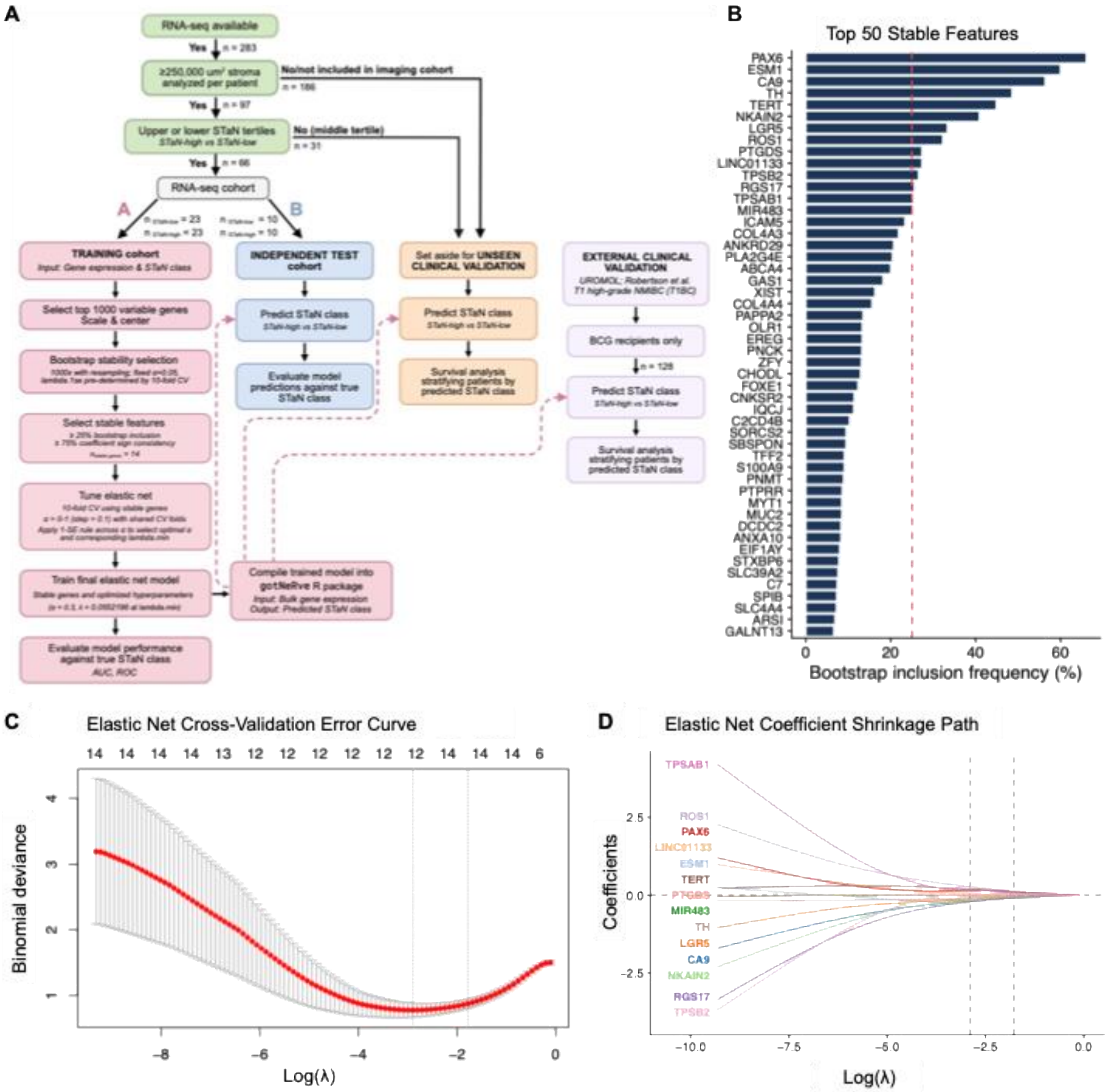


**Supplementary Figure 6. Development and validation of an elastic net classifier to predict STaN class (STaN-high or STaN-low) from bulk RNA sequencing. (A)** Workflow for development and validation of the classifier. The classifier and all associated code and data are available through gotNeRve R package at https://github.com/cdeit/gotNeRve. **(B)** Bootstrap feature selection frequencies for the top 50 genes across 1,000 bootstrap iterations. Bars indicate the percentage of iterations in which each gene was retained in the elastic net feature set. **(C)** Ten-fold cross-validation curve used to identify the optimal regularization parameter (λ). Red points indicate the mean cross-validation deviance, and gray error bars represent ±1 standard error. **(D)** Elastic net coefficient shrinkage paths showing changes in model coefficients as the regularization parameter (λ) increases.

**A**

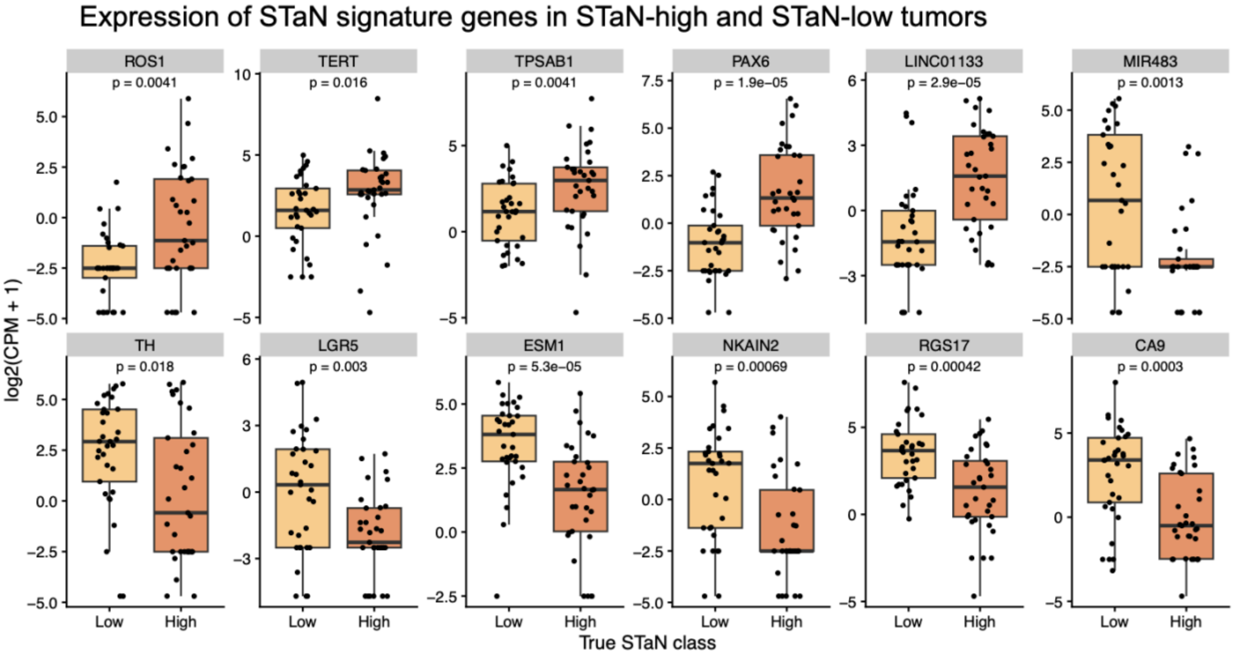


**Supplementary Figure 7. Expression of STaN 14-gene transcriptional signature genes in STaN-high vs STaN-low tumors. (A)** Normalized expression (log2 counts per million + 1) of genes comprising the STaN transcriptional signature in STaN-low (lower tertile; n = 36) and STaN-high (upper tertile; n = 36) tumors from the RNA-sequencing cohort. Boxes indicate the median and interquartile range, whiskers extend to 1.5 × IQR, and points represent individual tumors. Statistical significance was assessed using two-sided Wilcoxon rank-sum tests. **(B)** Volcano plot showing differential gene expression between STaN-high and STaN-low tumors. Differentially expressed genes were identified using DESeq2 with thresholds of $|\log_2(\text{fold change})| > 1$ and false discovery rate (FDR) < 0.05. Red and blue points represent genes significantly upregulated or downregulated in STaN-high, respectively. Gray points represent genes that are not significantly differentially expressed. Genes included in the STaN 14-gene transcriptional signature are labeled.

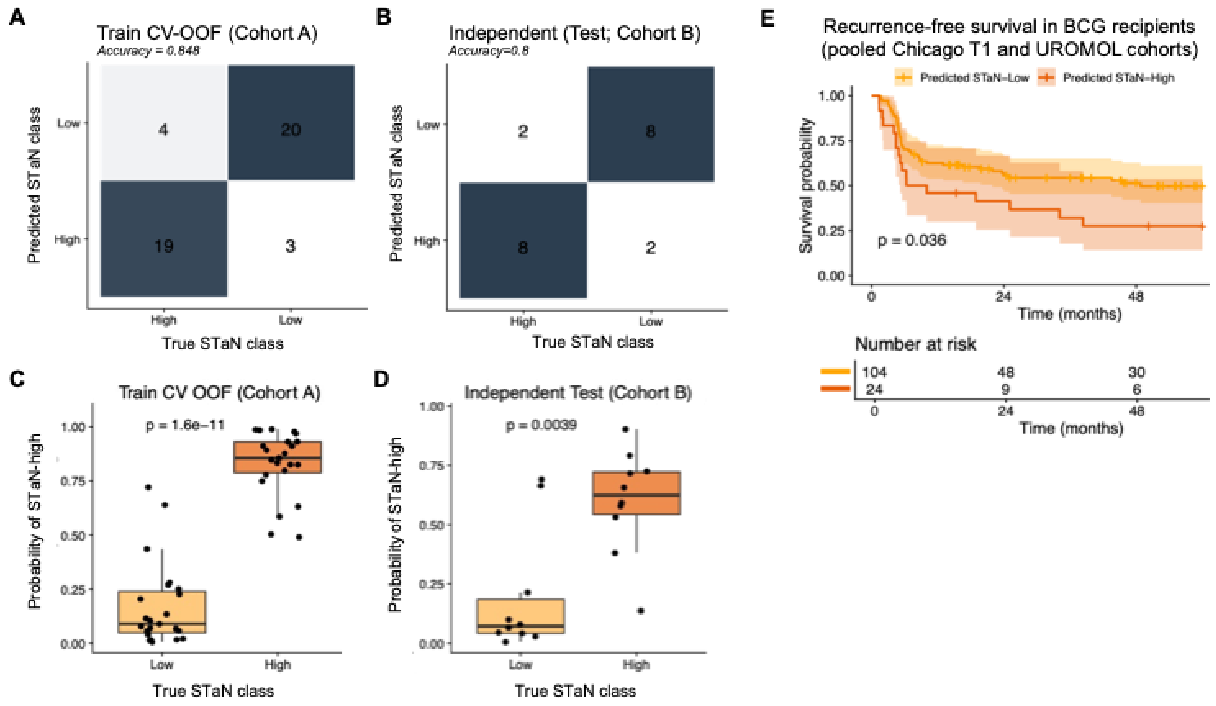


**Supplementary Figure 8. Performance of the STaN classifier and external clinical validation. (A,B)** Confusion matrices summarizing classifier performance in the training cohort using cross-validation out-of-fold predictions (CV OOF) **(A)** and the independent test cohort **(B)**. **(C,D)** Predicted probabilities of STaN-high classification stratified by true STaN class in the training cohort **(C)** and independent test cohort **(D)**. Boxes indicate the median and interquartile range (IQR), whiskers extend to 1.5x IQR, and points represent individual tumors. Statistical significance was assessed using two-sided Wilcoxon rank-sum tests. **(E)** Kaplan-Meier analysis of recurrence-free survival in BCG recipients from the combined external validation cohorts (Chicago T1BC and UROMOL), stratified by predicted STaN class. Shaded regions indicate 95% confidence intervals. Survival differences were assessed using the log-rank test.

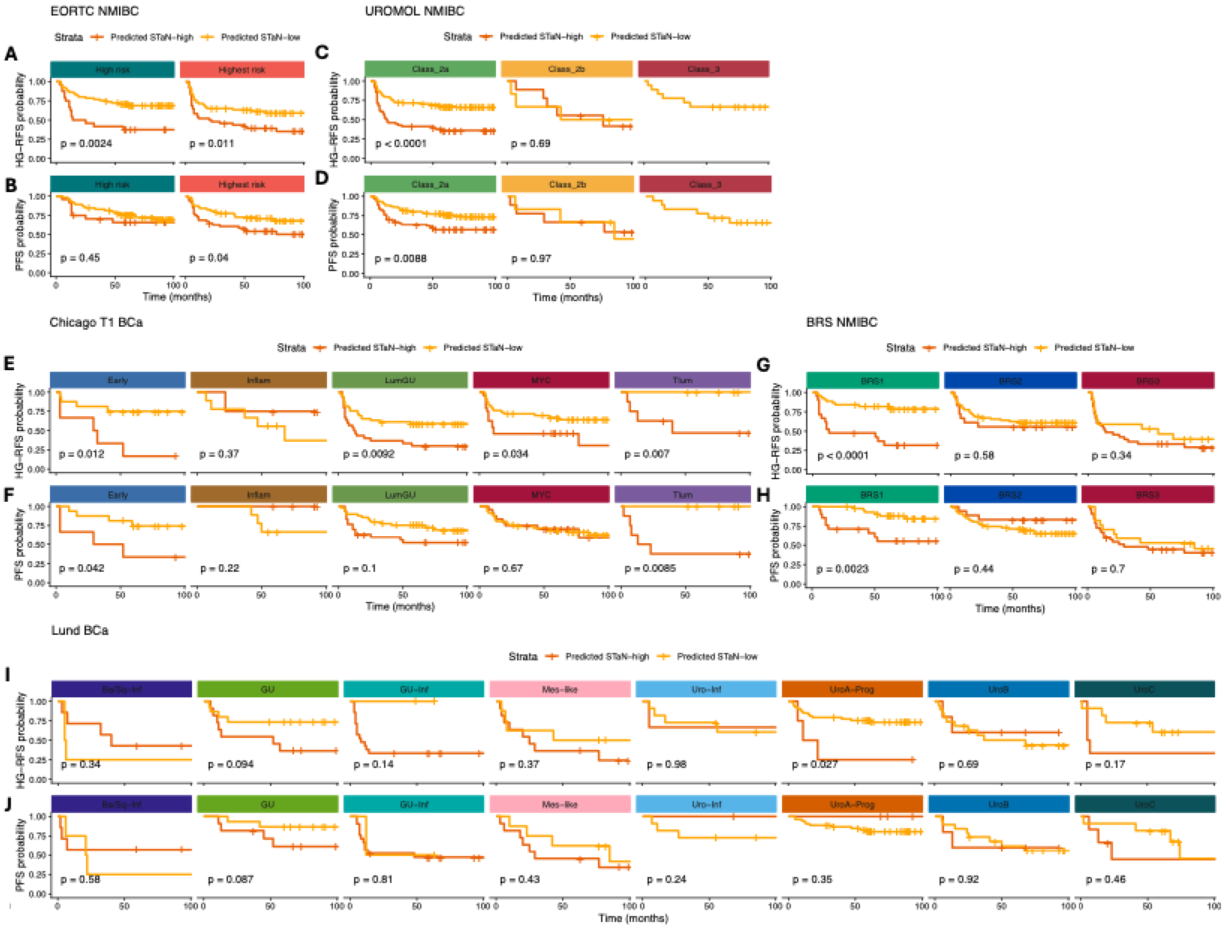


**Supplementary Figure 9. Predicted STaN class provides additional prognostic stratification within established NMIBC clinicopathological risk groups and molecular subtypes.** Kaplan-Meier analysis of high-grade recurrence-free survival (HG-RFS) and progression-free survival (PFS) stratified by predicted STaN abundance within established clinicopathological risk groups and molecular subtype classification systems in the unseen NMIBC cohort (n = 200). Patients were classified as predicted STaN-high or STaN-low using the gotNeRve transcriptomic classifier and subsequently evaluated within **(A,B)** EORTC, **(C,D)** UROMOL, **(E,F)** Chicago T1, **(G,H)** BRS, **(I,J)** and Lund molecular subtypes. Survival differences were assessed using the log-rank test. P-values are shown for each comparison.

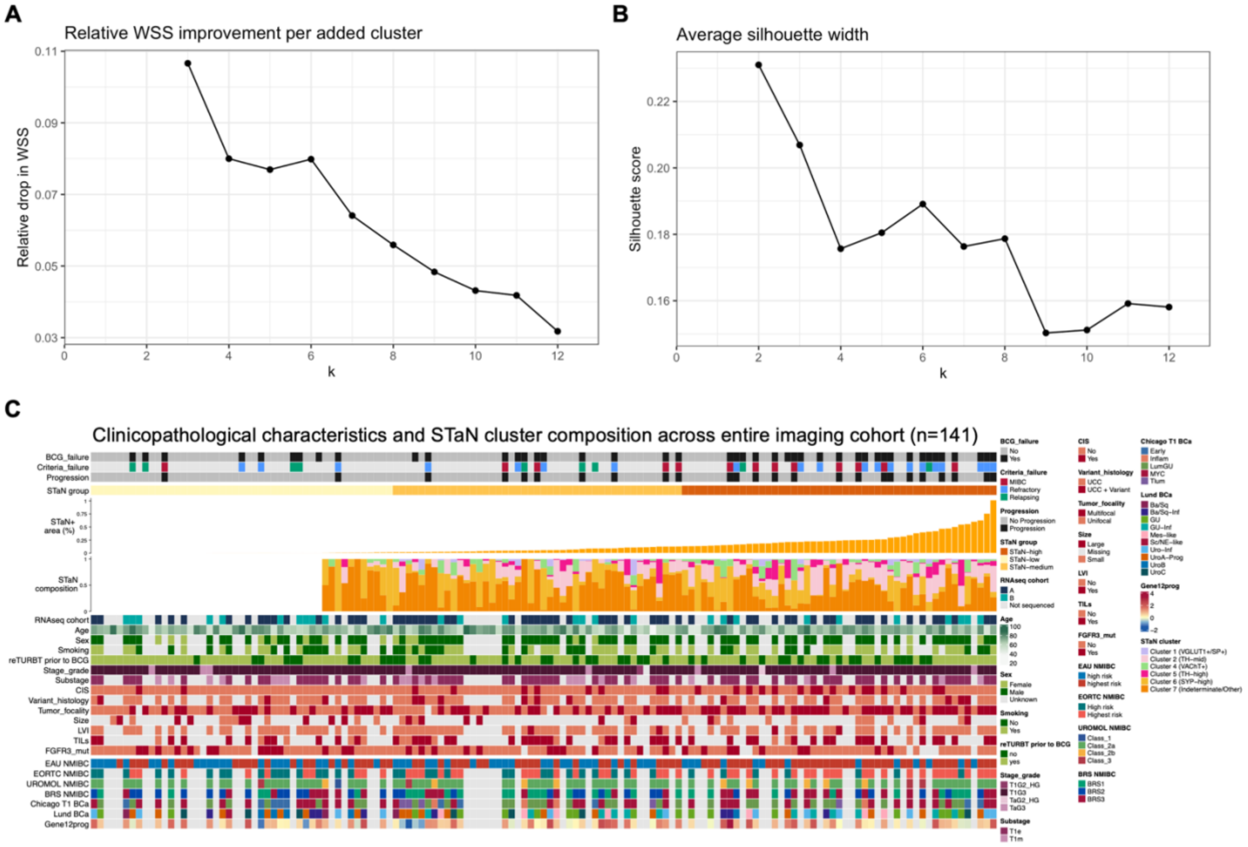


**Supplementary Figure 10.** STaN phenotype classification by k-means clustering. **(A,B)** Selection of the optimal number of clusters using the within-cluster sum of squares (WSS) method **(A)** and average silhouette score **(B)**. **(C)** Heatmap showing k-means cluster assignment derived from multiplex STaN marker expression. STaN abundance, nerve subtype composition, clinicopathological variables, and molecular features are displayed for comparison across the entire final imaging cohort (n = 141).
